# Capillary bundling of microtubules by condensates

**DOI:** 10.64898/2026.06.19.733462

**Authors:** Bernardo Gouveia, J. Pedro de Souza, Venecia Valdez, Joshua W. Shaevitz, Howard A. Stone, Sabine Petry

## Abstract

The cytoskeleton organizes the cellular interior using cytoskeletal filaments that rely on bundling, usually executed by stable and ordered crosslinking proteins. Bundling often requires protein complexes with at least two defined microtubule binding regions, as present in many molecular motors. Here, we establish a mechanism of microtubule bundling based on capillary forces, analogous to how wet hair sticks together. We show using *in vitro* experiments and theory that condensates can bundle microtubules through capillary forces, wherein liquid-like capillary bridges form between microtubules and adhere them together through interfacial and wetting forces. We quantify the structure and dynamics of these capillary bundles using total internal reflection fluorescence microscopy, and directly measure the charge-dependent interfacial tensions of condensates on microtubules using atomic force microscopy. Lastly, we show that these capillary bridges provide viscous resistance to motor-driven microtubule sliding that is insensitive to the bulk protein concentration. Taken together, we provide a novel mechanism for how cytoskeletal filaments bundle: through condensate-mediated capillary forces.

## I. INTRODUCTION

As part of the eukaryotic cytoskeleton, microtubules are rigid, tubular polymers that form the structural foundation of the mitotic spindle [1], the neuronal axon [2], and cilia [3]. Microtubules often form bundles with other microtubules to generate larger, more stable structures, such as kinetochore fibers [4, 5]. The standard view of how microtubule bundles form is through the action of crosslinking proteins that bridge two microtubules together through at least two definable microtubule binding regions that typically have positive charge [6–8]. Passive crosslinkers range from the physiological dimer PRC1 [9, 10] to small, synthetic peptides with multi-valent, positively charged residue strings [11]. Active crosslinkers include the tetrameric motor protein kinesin-5 (Eg5/KIF11) [12, 13] and the dimeric motor protein kinesin-14 (Ncd/HSET) [13, 14]. In these examples, the mechanism of microtubule bundling is readily apparent: these proteins form stable multimers or have multivalent motifs that can directly bind two microtubules simultaneously using multiple microtubule binding regions.

There has been recent interest in studying microtubule associated proteins (MAPs) that form biomolecular condensates above a saturation concentration—that is, MAPs that can thermodynamically partition into a dilute phase and dense phase (the condensate). Even though the MAP content of the condensate is elevated, it often exhibits liquid-like properties [15, 16]. Although the focus in the literature has been on how condensed MAPs regulate active microtubule processes such as dynamic instability and microtubule nucleation, it is striking that in nearly all of these studies microtubule bundling is also observed. Examples include TPX2 [17– 20], BuGZ [21], Tau [22–24], LEM2 [25], CLIP-170 [26– 28], MAP65 [29, 30], TPPP [31], and CPC components [32] both *in vitro* and *in vivo*. This is not only true for microtubules; actin-binding proteins that form condensates have also been shown to bundle actin filaments, with examples including VASP [33], the Nephrin– Nck–NWASP system [34, 35], and the LAT signaling system [36]. Thus, there seems to be a general relationship between the ability of cytoskeletal-associated proteins to form condensates and their ability to bundle filaments. While this relationship has been noticed in the literature [37], the precise mechanism by which this occurs remains to be determined.

Most of these MAPs are composed of large intrinsically disordered regions with only one microtubule binding domain, if at all. They form condensates through multivalent, weak interactions [16]. Due to the seemingly generic nature of these bundling observations, it is unlikely that condensed MAPs bundle microtubules by stably dimerizing and bridging two filaments together through multiple defined binding regions. We hypothesized that a more mesoscale mechanism must be at play operating above the length scale of single protein interfaces, especially in the vicinity of or above the saturation concentration for biomolecular condensation.

Specifically, we hypothesized that condensed MAPs can bundle microtubules through capillary forces, which occur when a liquid phase wets a solid substrate and result from the interfacial tension of the liquid-liquid interface [38]. These forces can create liquid capillary bridges between two substrates, and are precisely the everyday forces we experience when a wetted finger turns the page of a book, or when wet hair sticks together, and are common in millimeter-to-micrometer scale wetted fiber networks [39, 40]. Due to the liquid-like nature of biomolecular condensates and the variety of soft substrates available to wet, such as chromatin [41, 42], cytoskeletal filaments [43–45], and membranes [46, 47], interfacial forces are already being implicated in diverse intracellular contexts that involve condensates [48].

To test this bundling hypothesis, we employed total internal reflection fluorescence microscopy (TIRFM) and atomic force microscopy (AFM) experiments on model proteins TPX2 and BuGZ *in vitro* in combination with theoretical modeling. We show that condensed TPX2 and BuGZ can bundle microtubules via liquid-like capillary bridges, and that this mechanism is fundamentally distinct from bundling via single-molecule crosslinkers.

## II. RESULTS

### A. Condensed MAPs bundle microtubules through capillary bridges *in vitro*

To study how condensed MAPs bundle microtubules, we used purified recombinant BuGZ-BFP and GFP-TPX2 as our model proteins throughout this paper. Both have been reported previously to phase separate into condensates above a bulk saturation concentration [20, 21]. However, whereas BuGZ only has one microtubule binding region [21], TPX2 has two [49], and so in principle it could bundle microtubules through direct crosslinking as well as through condensation. We performed a bulk phase separation assay and found that BuGZ condensed at ≈140 nM while TPX2 condensed at ≈50 nM in assay buffer (Fig. S1, Supplementary Methods). We note that condensation on microtubules will occur at a concentration lower than the bulk saturation concentration due to prewetting effects [45, 50], so the bulk saturation value can only be taken as a rough guide.

Next, we mixed short (1–5 *µ*m in length, 25 nm in diameter), stabilized, ATTO647-labelled GMPCPP microtubules with either BuGZ or TPX2 at different concentrations and flowed the mixture into a flow chamber with the coverslip side down. After a 10 min incubation, bundles settled to the coverslip surface and were imaged with TIRFM. For BuGZ, robust microtubule bundles were only observed near and above the bulk phase boundary (Fig. 1a). For TPX2, microtubule bundling began between 12.5–25 nM (Fig. 1b). We quantified bundle formation with an order parameter defined as 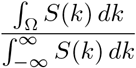, where *S*(*k*) is the Fourier-space structure factor of the TIRFM images and Ω is a set of modes that captures the bundling transition (Fig. 1c-d, Fig. S2, Supplementary Methods). For BuGZ, the bundling transition is clearly sigmoidal, consistent with dynamics driven by phase separation (Fig. 1c). The transition occurs just before the bulk phase boundary, likely due to prewetting effects. For TPX2, the bundling transition occurs over a larger concentration range, which may be attributed to TPX2’s ability to crosslink microtubules at the singlemolecule level [49] as well as bundle them through condensation (Fig. 1d). For both proteins, we note the high background signal of soluble tubulin below the phase boundaries. Above the phase boundary, soluble tubulin gets sequestered into condensed MAPs as previously reported [20, 21], increasing the intensity contrast of the microtubule channel.

**FIG. 1.**
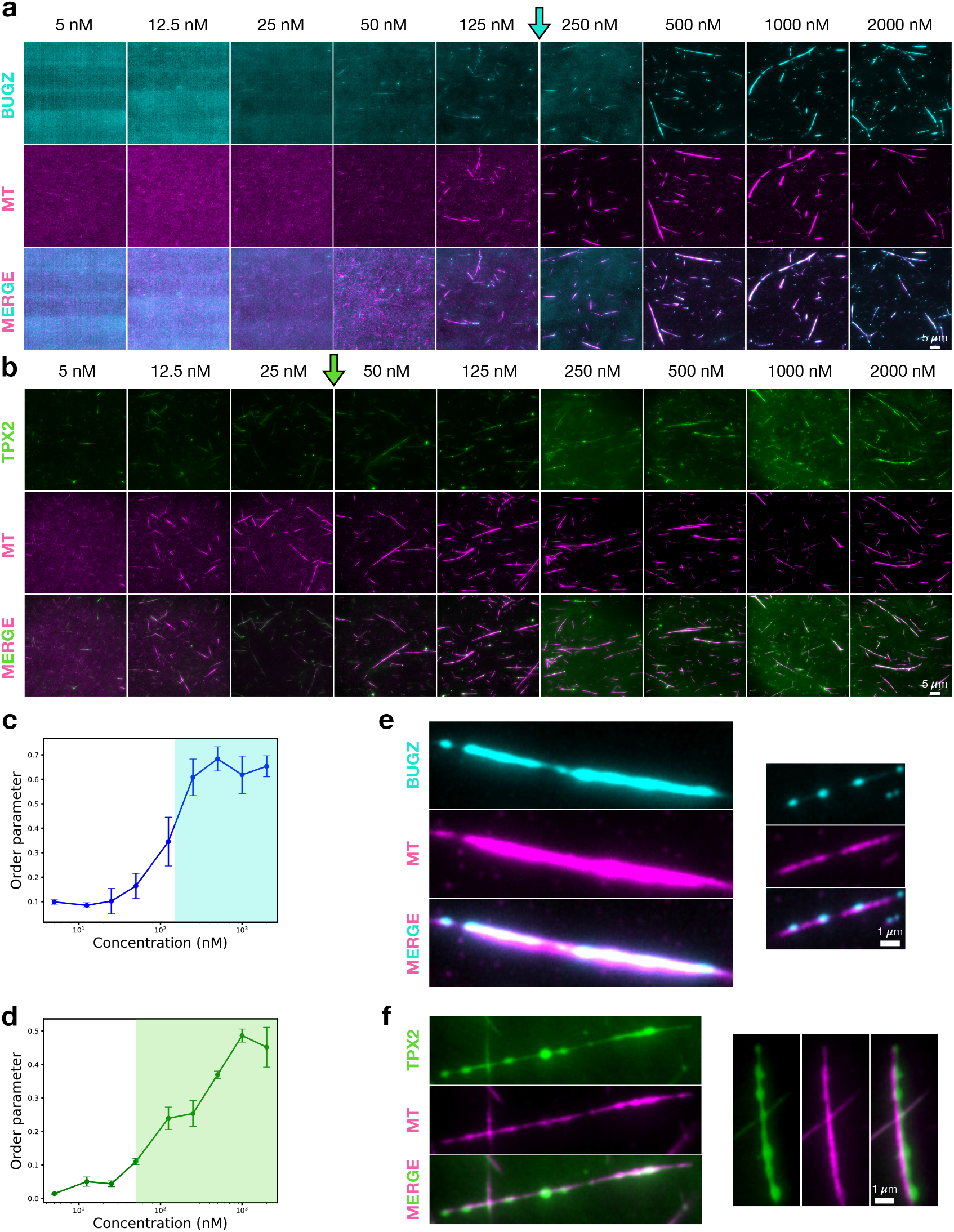
Condensed microtubule-associated proteins (MAPs) bundle microtubules. TIRFM images of microtubule bundles that sedimented to the flow channel surface after a 10 min incubation with **a**. BuGZ (1 microtubule binding region) and **b**. TPX2 (2 microtubule binding regions). Colored arrows indicate concentration above which these MAPs form condensates in bulk solution. Microtubule channel lookup tables are the same across concentrations to enable direct comparison. Note the decrease in background soluble tubulin above the phase boundaries, indicating partitioning into condensates. MAP channel look-up tables are optimized per concentration to allow for visualization. Scale bars are 5 *µ*m. Order parameter that describes the bundling transition for **c**. BuGZ and **d**. TPX2 as a function of bulk MAP concentration. The order parameter is calculated as 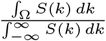, where *S*(*k*) is the Fourier-space structure factor and Ω is the set of modes that captures the bundling transition (Fig. S2). Error bars are standard deviations from *N* = 5 images per concentration. Shaded regions demarcate concentrations above the bulk phase boundaries. Zoomed in TIRFM images of **e**. BuGZ and **f**. TPX2 bundles both at 2000 nM displaying film and droplet-like structures expected for filaments held together by capillary forces. Scale bars are 1 *µ*m.

Liquid-like features, such as droplets and films, on microtubule bundles that are larger than the diffraction limit are visible for both BuGZ (Fig. 1e) and TPX2 (Fig. 1f) at sufficiently high MAP concentrations, since an increase in bulk MAP concentration above the phase boundary serves to increase the volume of the condensed phase [43]. Furthermore, decreasing the microtubule density in the assay by a factor of 10 resulted in smaller bundles for a given MAP concentration, but larger droplets along those bundles (Fig. S3). This observation is a consequence of more volume of condensed MAP per available microtubule surface area, resulting in bigger droplets along bundles [43]. A negative control in the absence of MAPs ruled out microtubule bundling from other forces such as depletion interactions [51] under our experimental conditions (Fig. S4). Such liquid-like features share striking resemblance to the micrometer-to-millimeter scale wetted fiber networks that are known to form bundles through capillary forces [39, 40].

As another control, we performed bundling experiments using Eg5 at low ATP concentrations (1 *µ*M), where it can still bind and crosslink microtubules but not slide them apart. We first confirmed that “static Eg5” does not phase separate (Fig. S5a), and thus can serve as a model of a bundler that acts solely by singlemolecule crosslinking. In the bundling assay, bundles started to settle on the coverslip at concentrations above 2.5 nM Eg5 (Fig. S5b). This is an order of magnitude lower than BuGZ (above 50 nM), which can only bundle microtubules by condensation, and is comparable to TPX2 (above 5 nM), which can bundle microtubules via both mechanisms. No droplet or film structures were visible for static Eg5 bundles at any concentration. Furthermore, we do not observe any sequestering of soluble tubulin into regions of high Eg5 intensity, like we did for the condensed MAPs. Taken together, these observations motivated us to quantitatively investigate whether capillary forces are the dominant bundling mechanism for condensed MAPs.

### B. Direct measurement of condensate-driven capillary forces on microtubules

We sought to directly measure the magnitude of these capillary forces on microtubule surfaces by using in-fluid AFM. GMPCPP stabilized microtubules were electrostatically adhered to an atomically smooth mica surface and either TPX2 or BuGZ was added to a final concentration above their bulk phase boundary value (Fig. 2a). Microtubules coated with condensed MAP were first located using fast-tapping-mode AFM, after which we performed a series of slow approach-retract force ramps at specified locations along the lengths of microtubules (Supplementary Methods). The average force curves (Fig. 2c-d) exhibit significant adhesion compared to measurements of naked microtubules in the same buffer (Fig. 2b), both in the retract and approach ramps, where the magnitude of the adhesive force is the minimum of the force curve (Figs. 2e-f). We interpret these adhesive forces as condensate-mediated capillary bridges between the microtubule and the AFM silicon-nitride tip, both of which are negatively charged in our experimental conditions [52].

**FIG. 2.**
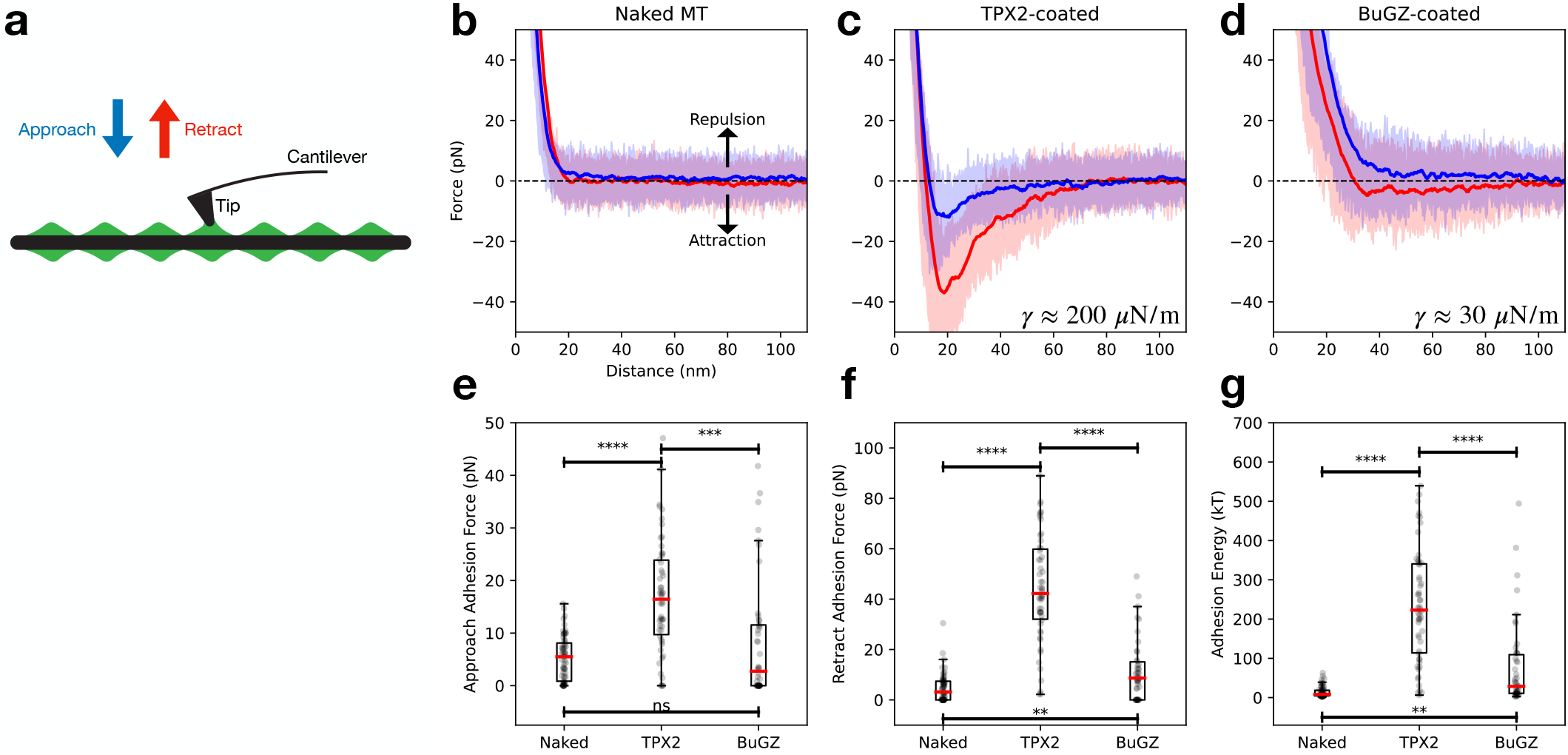
AFM measurements quantify interfacial forces. **a**. Schematic of experimental in-fluid AFM configuration. First, a microtubule is found using an AFM probe of tip radius *R*_*T*_ ≈ 20 nm. A series of approach/retract force spectroscopy curves are then instantiated along the coated microtubule. Average approach (blue) and retract (red) curves for **b**. a naked microtubule (*N* = 65), **c**. a TPX2-coated microtubule (*N* = 54), and **d**. a BuGZ-coated microtubule (*N* = 46). The shaded blue/red regions correspond to the 25th and 75th percentile of approach/retraction force curves. Statistics of the **e**. approach adhesion force, **f**. retract adhesion force, and **g**. adhesion energy for naked, TPX2-coated, and BuGZ-coated microtubules. The red line is the median. Interfacial tensions for condensed TPX2 and BuGZ were estimated using the formula 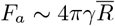, where *F*_*a*_ is the average retract adhesion force and 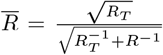 where *R*_*T*_ ≈ 20 nm is the nominal AFM tip radius and *R* ≈ 13 nm is the microtubule radius.

Using the minima of the averaged retract curve (Fig. 2c-d) as a measure for the adhesive force *F*_*a*_, we can estimate the interfacial tension of each condensate via 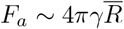, where the effective radius is 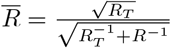 (Supplementary Theory). This force expression assumes a highly idealized contact configuration of the capillary bridge extent being small relative to the AFM tip radius and microtubule radius, yet it can be used as an estimate for calculating the surface tension. *R*_*T*_ ≈ 20 nm is the nominal AFM tip radius and *R* ≈ 13 nm is the microtubule radius. We find *γ*_TPX2_ ≈ 200 *µ*N/m and *γ*_BuGZ_ ≈ 30 *µ*N/m. These interfacial tension values measured with a nanoscale probe fall in the higher end of the range of previous measurements of condensate interfacial tensions that relied on other techniques that used micron-scale probes [53]. Indeed, there are subtle differences arising at the nanoscale, such as the relative importance of the interface thickness [44], that could influence the effective measured value of *γ*. By measuring these forces at the nanoscale (Fig. 2b-f), we obtain a direct estimate of the relevant forces protein complexes and intracellular surfaces would experience from a single capillary bridge on microtubules.

To explain the order of magnitude difference in interfacial tensions between condensed TPX2 and BuGZ, we start with a Flory–Huggins-type phase field model [54– 56] (Supplementary Theory). We find that the interfacial tension scales like

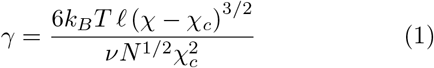

near the critical point, where *χ* is the mean-field interaction parameter, 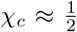, *N* is the degree of polymerization, and *ℓ* and *ν* are the length and volume, respectively, that characterize the lattice size in the theory. The surface tension expression in Eq. (1) is derived only for regions in the phase diagram near the critical point, and may only be used approximately when relating the protein characteristics to their condensed phase surface tension more generally. Because *N* is similar between TPX2 and BuGZ, the difference in *γ* likely arises due to a difference in molecular interaction strength *χ* that drives phase separation. If we assume such interactions are dominated by electrostatics, a modified Voorn–Overbeek theory [57–59] gives *χ* ∼ *f* ^2^, where *f* is the fraction of charged amino acids in the protein chain (Supplementary Theory). We note that the Voorn–Overbeek theory applied here accounts for the effective electrostatic free energy of both positive and negatively charged sites along the polyampholytic protein backbone. While the predictions are limited by standard mean-field assumptions, the model can provide a useful estimate of the strength of the electrostatic interactions driving phase separation. We estimate *f*_TPX2_ ≈ 0.33 and *f*_BuGZ_ ≈ 0.20, which leads to *χ*_TPX2_≈ 2.3 and *χ*_BuGZ_ ≈ 0.9. This results in a prediction of 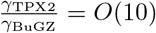, which rationalizes the measured difference in *γ*. Thus, our theory connects sequence charge features of proteins to the interfacial tension of their condensed phase.

The AFM force curves can be integrated to generate energy curves, the minimum of which is the capillary adhesion energy *E*_*a*_. We measure adhesion energies per unit area 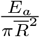 in the range of 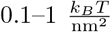 (Fig. 2g). We note that these adhesion energies are at least an order of magnitude higher than those measured for tubulin dimers on the interfaces of stress granules [44]. While this might reflect that condensed MAPs offer stronger adhesion specifically to tubulin lattices, we caution that comparing *in vitro* material properties to those *in vivo* is a qualitative endeavor, since condensates can be in different parts of their phase diagrams depending on environmental context, which affects the partition coefficient and therefore the interfacial tension [60].

In addition, we leveraged our AFM technique to measure the viscosities *µ* of the condensates (Fig. S6). To do so, we let bulk MAP droplets (sizes ≳ 1 *µ*m) sediment to the glass bottom of a dish and performed a force ramp with a larger 1 *µ*m nominal tip radius (Supplementary Methods). Using a lubrication analysis, we find average values of *µ*_TPX2_ ≈ *µ*_BuGZ_ ≈ 500 mPa s for the condensed MAPs and *µ*_*s*_ ≈ 1 mPa s for the assay buffer. We note that the condensates get more viscous over the timescale of minutes as they age, in agreement with previous observations (Fig. S6f-g) [61].

### C. Snapping dynamics of capillary bundles

To rationalize the dynamics of bundle formation, we developed a theoretical model where two microtubules contact each other at an arbitrary initial angle *ψ*_0_ that evolves in time as *ψ*(*t*) via a liquid capillary bridge with contact angle *θ* (Fig. 3a). There is both an adhesive capillary force [62] that acts to bring the microtubules closer together and a capillary torque [63, 64] that acts to drive the liquid-liquid interface to a state of minimal energy as well as to maximize the contact the condensate has with the microtubules. We assume that the condensate interface relaxes more quickly than the rate of microtubule motion, so that it is always locally equilibrated (Supplementary Theory). This predicts an equilibrium state of bundled microtubules that are parallel and wetted by the condensate.

**FIG. 3.**
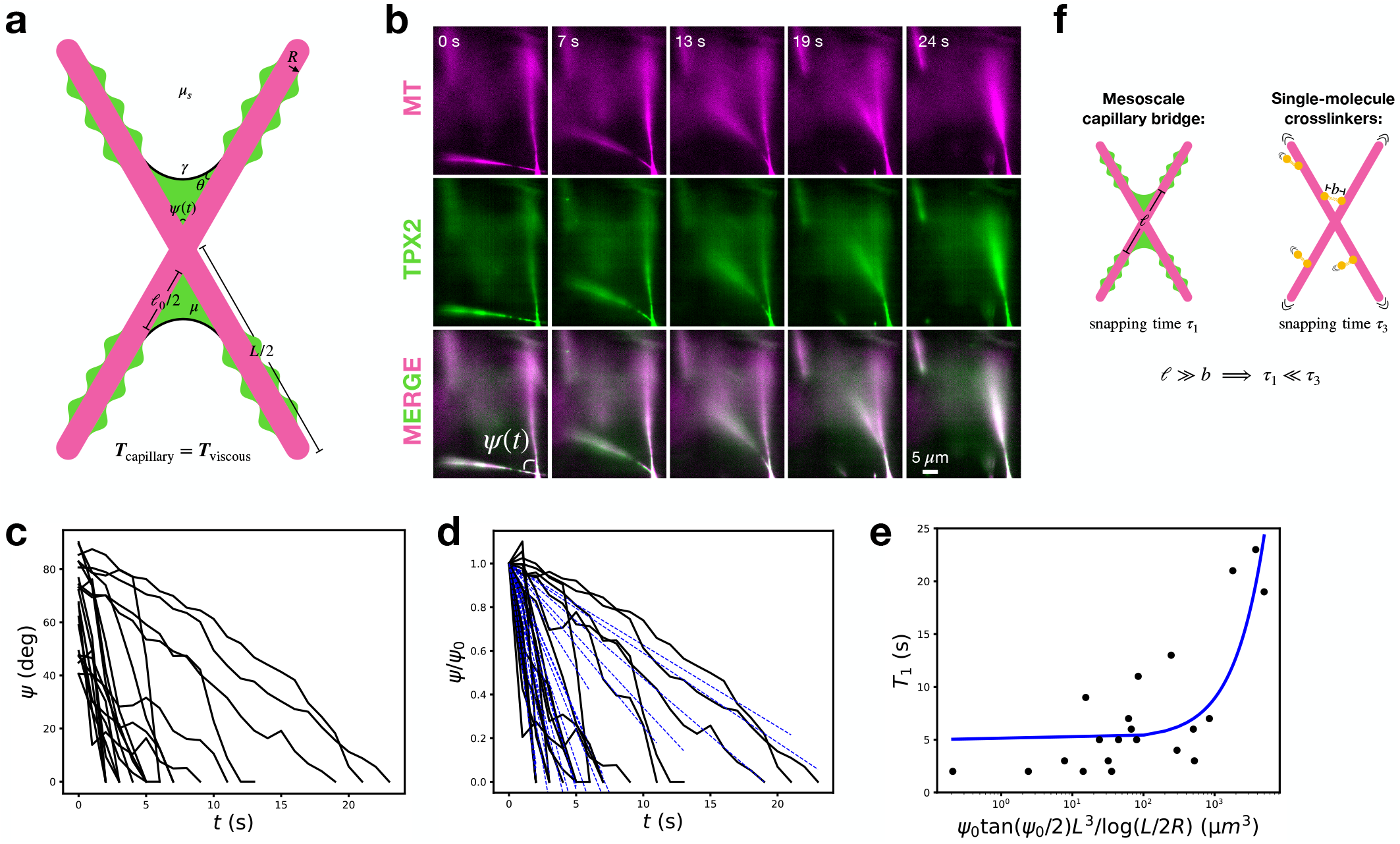
Snapping dynamics of capillary bundles. **a**. Schematic showing the non-equilibrium configuration of two microtubules of length *L* that contact at an arbitrary angle *ψ*(*t*). The microtubules are wetted by a condensate with surface tension *γ*, viscosity *µ*, and contact angle *θ* in a solvent of viscosity *µ*_*s*_. **b**. Oblique TIRFM live imaging of two bulk microtubule bundles coated with condensed TPX2 above the phase boundary (500 nM) snapping together. Background subtraction was done to enhance visual contrast. Scale bar is 5 *µ*m. **c**. Plot of oblique TIRFM measurements of snapping angles versus time. *N* = 21 snapping events. All data is taken with TPX2 at 500 nM. **d**. Plot of snapping angles versus time overlaying least-squares fits to Eq. (2) (dashed blue lines). **e**. Plot of snapping times versus the length of the shortest microtubule in the pair. The blue curve is the least-squares fit to the function 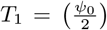 tan 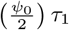, where 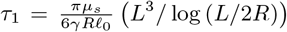 log (*L*^3^*/*log 2*R*). All parameters are independently measured or known except *ℓ*_0_, which we use as a least-squares fitting parameter, finding *ℓ*_0_ = 9 *±* 1 nm. **f**. Schematic contrasting capillary bundling versus single-molecule crosslinking. Because the wetted length *ℓ* is a mesoscale length scale that can approach the microtubule length *L* for late snapping times, we expect capillary torques to be generically larger and snap microtubules more quickly than crosslinking torques, whose moment arm is limited by a molecular length scale *b*.

For early times (*ψ* → *ψ*_0_) the capillary torque is primarily resisted by dragging the filaments together through the solvent of viscosity *µ*_*s*_. Assuming perfect wetting (*θ* = 0) we find that the angle *ψ* should decrease linearly in time as

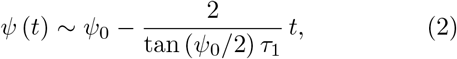

where 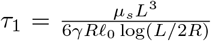, *L* is the microtubule length, *R* is the microtubule radius, *γ* is the interfacial tension, and *ℓ*_0_ is the initial wetted length of the droplet that contacts both filaments.

To test these ideas, we mixed short, stabilized, ATTO647-labelled GMPCPP microtubules with 500 nM TPX2 in a well and imaged bundle formation live in 3D using oblique TIRFM (Supplementary Methods). We directly observed TPX2-coated microtubules contacting at an initial angle and snapping together until they formed a parallel bundle, as predicted (Fig. 3b, Fig. S7). We also quantified individual snapping trajectories versus time (Fig. 3c-d), finding linear behavior at our experimentally observable temporal resolution, validating Eq. (2).

Using Eq. (2), we can compute the time it takes mic_(_rotu_)_bules_(_ to _)_fully snap together as 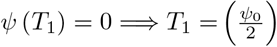 tan 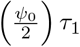. We fit this model to our data, where the only fit parameter is *ℓ*_0_, since we measured *µ*_*s*_ and *γ* using AFM and take *R* = 12.5 nm. The model captures the data reasonably well across our entire range of measured *L* and *ψ*_0_ (Fig. 3e), where for *L* we take the smallest microtubule length in the pair. From the fitting we learn *ℓ*_0_ = 9 *±* 1 nm, which is on the order of *R*, which is geometrically sensible for the initial wetted length upon microtubule contact.

How does this mechanism of microtubule bundling via capillary bridges fundamentally differ from bundling via single-molecule crosslinkers? During the late time (*ψ* → 0) dynamics of capillary snap✓ping, the wetted length increases significantly as 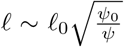 and the angle decreases as 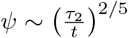 (Supplementary Theory). In this regime, the squeezing of the intervening condensate dominates the viscous torque at shallow angles. The governing time scale 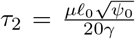 results from the balance of the capillary torque and the viscous resistance from squeezing the condensed film of viscosity *µ* between aligning filaments. Measuring these late time dynamics offer a way of probing a feature unique to capillary bridges. However, for the parameters we measure *τ*_2_ = *O*(10^−5^) s, which is too fast to observe using our current experimental set up.

In standard models of microtubule bundling by single-molecule crosslinkers, the crosslinkers are modeled as molecular springs of a characteristic length *b* that can bind two microtubules [7, 8]. Such a model predicts a characteristic time scale 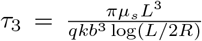, where *k* is the spring constant and *q* is the crosslinker density per unit microtubule length (Supplementary Theory). If we assume similar adhesion force scales between the two models so *kb* ∼ *γR*, and we take *qb* ∼ 1, we find that the ratio of timescales is given by 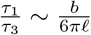. Because *b* is a molecular length scale and *ℓ* is the mesoscopic length scale of the wetted film, we expect *τ*_1_ ≪ *τ*_3_ (Fig. 3f). That is, even for comparable adhesion forces, we generally expect capillary torques to be at least an order of magnitude larger than crosslinking torques due to their mesoscale nature. For example, in the study of synthetic microtubules crosslinkers by Drechsler et al. [11], it takes their microtubules ≈ 6 min to snap together, whereas all our snapping events occur in less than 30 s, which supports our bundling argument.

### D. Sliding dynamics of capillary bundles

In addition to the adhesive force that bundles microtubules together, we predict that capillary bridges should provide *viscous* resistance to intrabundle microtubule sliding. Relative microtubule sliding, usually driven by molecular motors, is prevalent in nature with examples ranging from mitotic spindle maintenance [65, 66] to paradigmatic active matter models [67]. Understanding the resistance provided to motor-driven sliding by condensate-mediated capillary bridges is therefore an important biophysical question, which will additionally yield further insights into how capillary bundles differ from microtubules bundled by single-molecule crosslinkers.

To assess this, we performed *in vitro* sliding assays using purified recombinant Eg5-GFP and monitored how motor-driven sliding velocities changed as we titrated in BuGZ from nanomolar concentrations to above the phase boundary (Fig. 4a). We focused on BuGZ to avoid a known binding interaction between TPX2 and Eg5 [68]. We first flowed in long (*>* 5 *µ*m) Alexa568-labeled, biotinylated GMPCPP microtubules into a flow chamber with a passivated and neutravidin-functionalized coverslip. We then serially flowed in 100 nM of Eg5, short ATTO647-labelled GMPCPP microtubules, and a final mixture of 100 nM Eg5 and the final bulk concentration *c*_0_ of BuGZ. The reaction was imaged using TIRFM and the Eg5-driven sliding speeds *V* of the short microtubules were measured by constructing kymographs for each mobile microtubule. In the absence of BuGZ (*c*_0_ = 0), we measured *V* = 1.8 *±* 0.4 *µ*m/min for Eg5-driven sliding, in excellent agreement with previous measurements (Fig. 4b) [69]. As we increased the concentration *c*_0_ of BuGZ, we observed two regimes of behavior. Below the phase boundary and prewetting transition (as defined by the onset of microtubule bundling in Fig. 1c), the sliding speed *V* decreased with increasing *c*_0_. However, above the prewetting transition we observe a saturation of *V* to a constant value with no further dependence on *c*_0_ (Fig. 4c-d, Fig. S8).

**FIG. 4.**
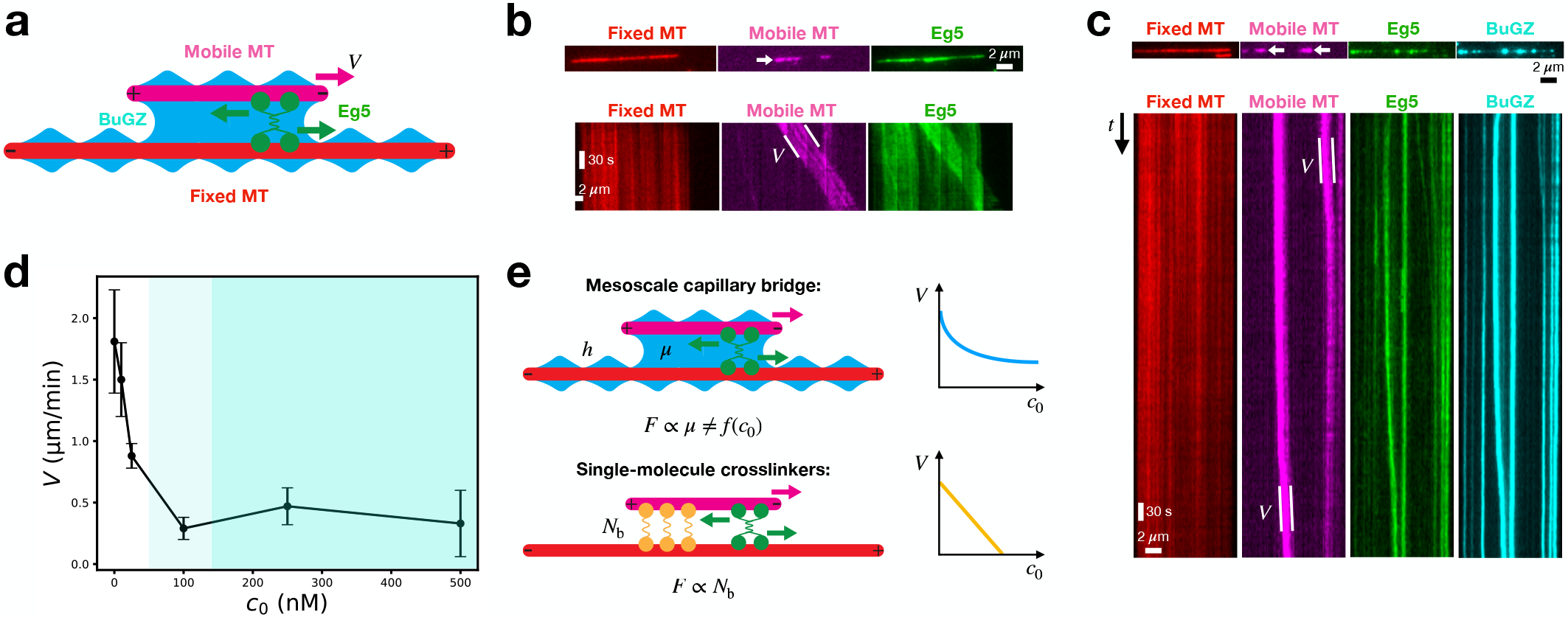
Sliding dynamics of capillary bundles. **a**. Schematic of the 4-color TIRFM experimental set up. Long, Alexa568-labeled biotinylated microtubules are strongly adhered to a passivated coverslip functionalized with neutravidin. Eg5 binds to these long microtubules and forms sandwiches with short, mobile ATTO647-labeled microtubules. If microtubules in the sandwich are antiparallel, Eg5 will slide the short, mobile microtubule towards its minus-end. The experiment is done as a function of BuGZ concentration below and above the bulk phase boundary. Snapshot and kymograph of a **b**. 0 nM BuGZ and 100 nM Eg5 experiment and a **c**. 500 nM BuGZ and 100 nM Eg5 experiment. The velocity *V* of the short, mobile microtubule is computed from the slope of the mobile microtubule kymograph. Scale bars are 2 *µ*m and 30 sec. **d**. Behavior of the sliding velocity *V* of the short, mobile microtubule as a function of the concentration of BuGZ *c*_0_ (black line). Error bars are standard deviations. *N >* 20 microtubules per condition. The darker shaded blue region represents BuGZ concentrations above the bulk phase boundary while the lighter shaded blue region is an estimate for the prewetting zone for BuGZ condensation based on the bundling transition in Fig. 1e. Below the phase boundary, *V* decreases with *c*_0_, whereas above the phase boundary, a viscous capillary bridge is established whose sliding resistance is insensitive to *c*_0_. Orange dashed line shows the predicted behavior if sliding resistance was due to single molecule cross-linkers that bind strongly enough to stall Eg5 sliding. **e**. Schematic comparing how single-molecule crosslinkers versus capillary bridges resist microtubule sliding. Crosslinkers bind and resist sliding in a manner proportional to the number of bound crosslinkers *N*_*b*_ until microtubules stall [70]. Near and above the phase boundary, condensates with a viscosity *µ* wet microtubules providing viscous resistance to sliding. Any further increase in bulk MAP concentration does not change the concentration of the dense phase nor its viscosity, and hence sliding resistance can saturate below the stall force.

We rationalize these observations as follows. Below the phase boundary, BuGZ binds the microtubule lattice at the single-molecule level and offers discrete roadblocks that can impair Eg5 sliding processivity (Fig. S8a-b). As *c*_0_ increases, more BuGZ binds, increasing the density of roadblocks and thus decreasing *V*. Near and above the phase boundary, condensed BuGZ fully wets the entire microtubule lattice and forms a capillary bridge with the mobile microtubule (Fig. 4c, Fig. S8c-d). In this scenario, the main resistance to sliding comes from the viscosity of the dense phase which scales like 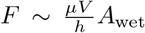, where *h* is the distance between microtubules and *A*_wet_ is the total area of microtubule lattice wetted by the capillary bridge. Since both microtubules are already fully wetted at this point, any further increase in *c*_0_ serves only to change the volume and shape of the bridge [62], which has a sub-dominant effect on *F*, as the majority of viscous shearing happens in the gap between microtubules. The crucial point here is that the condensate viscosity *µ* dominates the resistance, which is not a function of *c*_0_ above the phase boundary since the dense phase concentration remains constant (Fig. 4e).

How can we determine whether our measurements of condensed BuGZ viscosity are consistent with the decreased sliding velocity we observe in Fig. 4d? Using a simple model (Supplementary Theory), we find that the ratio of sliding velocities *V*_*f*_ */V*_*s*_, where *V*_*s*_ is the velocity with just buffer (viscosity *µ*_*s*_) between the microtubules and *V*_*f*_ is the velocity with an additional condensed film (viscosity *µ*) of thickness *h* between the microtubules, is

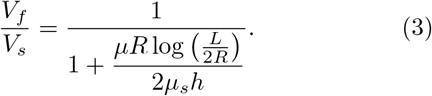

Keeping discussion at the level of order-of-magnitude estimates due to the measured spread and time-dependence of the viscosity (Fig. S6f-g), we have 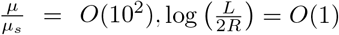, and 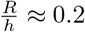, where we take *h* = 60 nm, which is the length of Eg5 [69]. This gives *V*_*f*_ */V*_*s*_ ≈ 0.1 which compares well to our measured value of 0.2, which we compute from Fig. 4d by averaging the velocities for 100–500 nM BuGZ and dividing it by the velocity for 0 nM BuGZ.

An alternative hypothesis is that bulk BuGZ condensates could sequester Eg5 into an inactive state, lowering the active population on the microtubule lattice and hence potentially lowering *V*. We ruled this out by performing sliding experiments at lower concentrations of Eg5, showing that *V* is insensitive to decreasing Eg5 concentrations over an order of magnitude (Fig. S9a-d). We also confirmed that while Eg5 density along microtubules measurably decreases upon first addition of BuGZ, it remains the same for all higher BuGZ concentrations (Fig. S9e). Taken together, the picture of constant Eg5 forcing across all BuGZ concentrations is reasonable, and thus we can attribute the decrease in sliding speed predominately to viscous resistance imparted by BuGZ.

This viscous resistance picture differs significantly from the sliding resistance imparted by crosslinkers that can individually bundle microtubules at the single-molecule level (Fig. 4e). For example, a similar assay [10] showed that the Eg5-driven sliding velocity *V* decreased in the presence of the homodimeric crosslinker PRC1 from 1.4 *±* 0.4 *µ*m/min at [PRC1] = 0.034 nM to 0.58 *±* 0.18 *µ*m/min at [PRC1] = 0.54 nM. This ∼ 60% decrease in *V* even at *<* 1 nM concentrations is in a totally different regime than our data (Fig. 4d), and reflects the fact that single-molecule crosslinkers provide sliding resistance at far lower concentrations than condensed MAPs that only bundle microtubules through viscous capillary bridges. Separate measurements using optical tweezers found that PRC1 crosslinkers resisted microtubule sliding in a manner directly proportional to the number of bound PRC1 crosslinkers [70], eventually resulting in microtubule stalling, which is different from the nonlinear response we measure here. Therefore, viscous capillary bridges can provide a way of generating resistance to motor-driven microtubule sliding that is below the motor stall force and insensitive to the bulk bundler concentration.

## III. DISCUSSION

In this work, we established that condensed MAPs can bundle microtubules through capillary forces. Using BuGZ and TPX2 as model MAPs, we observed robust microtubule bundling *in vitro* with droplet and film-like structures reminiscent of liquid capillary bridges optically visible above the bulk phase boundary. Using AFM, we directly measured the interfacial tension and viscosity of condensed MAPs, and derived a theory for how the interfacial tension depends on amino acid sequence charge. We then measured the snapping dynamics of live bundle formation and found agreement with a theoretical model of a capillary bridge snapping filaments together via the interfacial tension of the condensate. Lastly, we showed how Eg5-driven microtubule sliding is resisted by the presence of a condensed MAP, finding that above the phase boundary the resistance is insensitive to the MAP concentration. We hypothesize this is due to capillary bridges providing purely viscous resistance to sliding, in which the condensate viscosity is insensitive to the MAP concentration. Taken together, we presented a novel microtubule bundling mechanism, which is distinct from the previously established way of bundling via single molecule crosslinkers with at least two microtubule binding regions. Our results have several implications, which we discuss below.

First, *in vitro* observations of microtubule bundling by MAPs do not imply that the MAP can crosslink microtubules at the single-molecule level via two microtubule binding regions. A simple bulk phase separation assay that tests whether the MAP in question forms condensates at the experimental conditions of interest should be done before making any conclusions about the underlying microtubule bundling mechanism.

We stress that our model of microtubule bundling by capillary forces is insensitive to the molecular details that give rise to the condensed phase. The molecular nature of the interactions that give rise to condensates is an issue that is still debated [71, 72]. However, irrespective of the molecular mechanism of condensate biogenesis, as long as such a mesoscale compartment *exists* with internal fluidlike turnover kinetics and affinity for the microtubule lattice, capillary forces can be the dominant mechanism of microtubule bundling. Moreover, our usage of BuGZ and TPX2 in this work was purely as model proteins to study the biophysical aspects of capillary bundling. We do not claim that this mechanism is at play in living cells for these proteins, as that is likely context dependent and would require additional direct investigation [16].

While capillary bundling is a general mechanism that can explain how condensed MAPs bundle microtubules, other effects could be at play that do not rely on interfacial tension, especially between the prewetting transition and the bulk phase boundary. For example, the mechanism of tau-mediated microtubule bundling is complex and still debated, with hypotheses ranging from transient electrostatic dimers [73] to a viscoelastic network of crosslinked tau-tubulin oligomers [74]. While it is clear that condensed tau can form capillary bridges between microtubules under certain conditions [24], it likely depends on tau concentration, ionic strength, and other physicochemical factors. Even for the model proteins studied here, future separation of function mutants that can bind microtubules but not form condensates, and vice versa, would be helpful in truly isolating the effects of capillary forces in bundling by eliminating potential cooperative binding effects that might contribute to the bundling dynamics.

Our model assumes the simplest condensate rheology: that of a viscous Newtonian fluid. While this is sufficient to capture the main interfacial driving force, it is known that condensates in general are viscoelastic with frequency-dependent material properties [61, 75, 76]. How such rheology affects the dynamics of interfacialtension-driven processes such as capillary bundling is an area ripe for future research.

Because all our experiments are done under well-mixed conditions, the entire microtubule surface is coated with condensed MAP above the phase boundary. This is not necessary for the capillary bundling mechanism to operate. Rather, a condensed droplet could nucleate locally along any part of the microtubule, which can then go on to form a capillary bridge with another microtubule, thus utilizing protein amounts much more efficiently. This approach has already been employed to apply local capillary forces to genetic loci using condensates [42]. In the other extreme, over-expression of MAPs that can condense could lead to uncontrolled and unregulated microtubule bundling, which has already been observed for CLIP-170 [28] and TPX2 [77]. This concept could help explain certain cancer cell phenotypes [78] or other deleterious cell states.

The concept of bundling via capillary forces has been recently applied to rationalize the bundling behavior of actin filaments in the presence of FUS [79]. We anticipate that this mechanism will be relevant for intermediate filaments and septins as well. More broadly, the regulation and misregulation of condensates applying capillary forces on cytoskeletal filaments represent a fascinating direction for future study at the intersection of molecular biology, soft matter physics, and disease physiology [48].

## ACKNOWLEDGMENTS

We thank Sagar Setru, Yoonji Kim, Collin McManus, Sophie Travis, and Clifford Brangwynne for helpful discussions. B.G. was funded by the PD Soros fellowship, the NSF GRFP DGE-2039656, and the Wallace Memorial Honorific Fellowship. J.P.D. is supported by the Gong *91 Bioengineering Postdoctoral Fellows Program. This work was funded by NIH NIA 1DP2GM123493, Pew Scholars Program 00027340, Packard Foundation 201440376, and CPBF NSF PHY-1734030, and the Princeton Center for Complex Materials, a MRSEC (NSF DMR-2011750).

## AUTHOR CONTRIBUTIONS

B.G. performed theoretical analysis and experiments. J.P.S. performed theoretical analysis and experiments. V.V. designed and expressed the Eg5 construct and assisted with motor sliding experiments. H.A.S. performed theoretical analysis. J.W.S., H.A.S., and S.P. contributed to research design and mentoring. All authors wrote and edited the manuscript.

## ETHICS

Animal care was done in accordance with recommendations in the Guide for the Care and Use of Laboratory Animals of the NIH and the approved Institutional Animal Care and Use Committee (IACUC) protocol 1941-16 of Princeton University. Data and code used are available upon request.

## SUPPLEMENTARY FIGURES

**FIG. S1.**
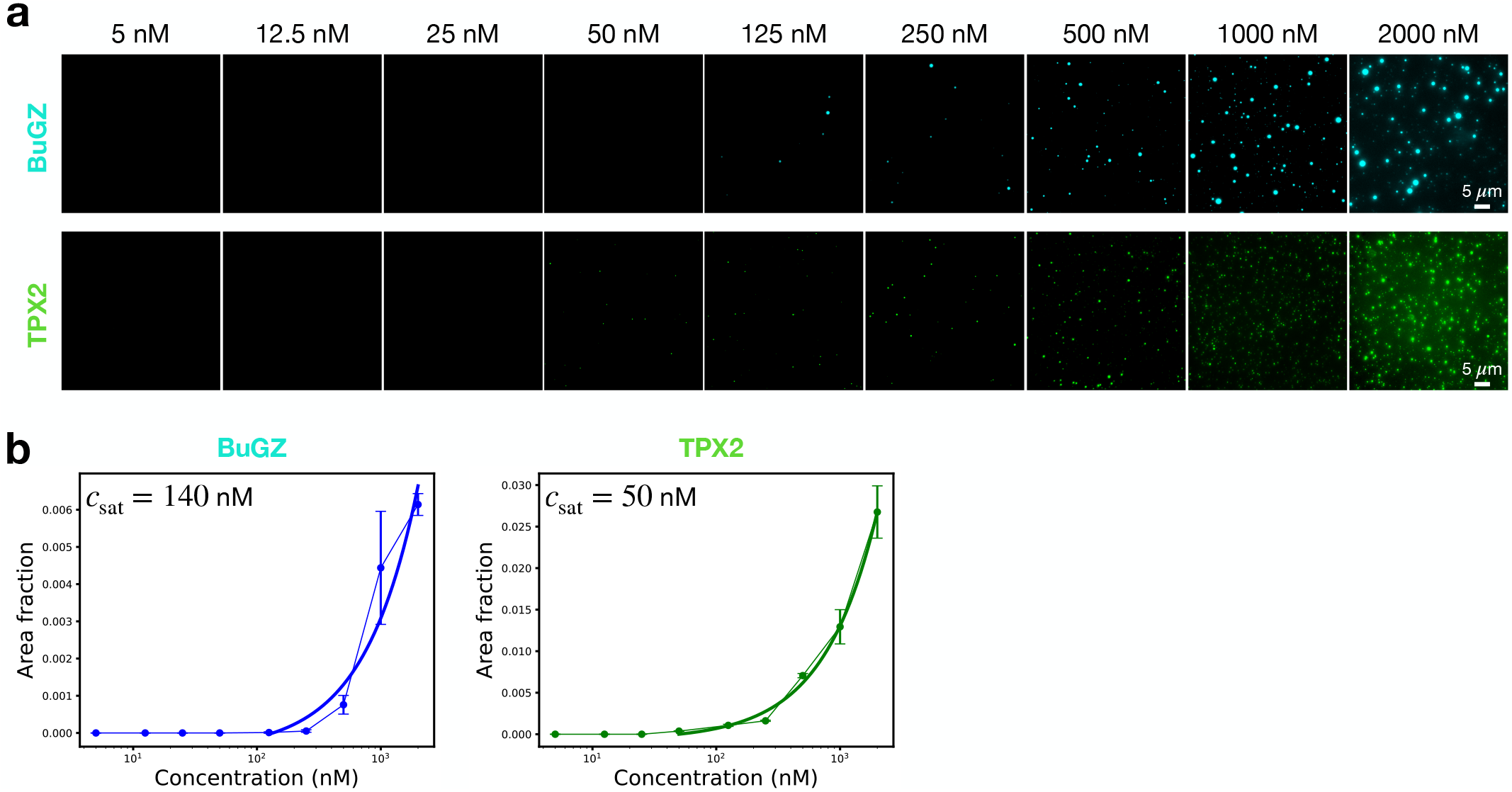
Bulk phase diagrams of BuGZ and TPX2. **a**. Epifluorescence images of bulk phase separation assays for BuGZ and TPX2. Lookup tables are the same across concentrations for each protein to enable direct comparison. Scale bars are 5 *µ*m. **b**. Phase diagrams for BuGZ and TPX2 showing area fraction of the condensed phase versus bulk concentration. The condensed phase was defined as regions having an intensity value above the threshold calculated using Otsu’s method for the highest concentration images for each protein.

**FIG. S2.**
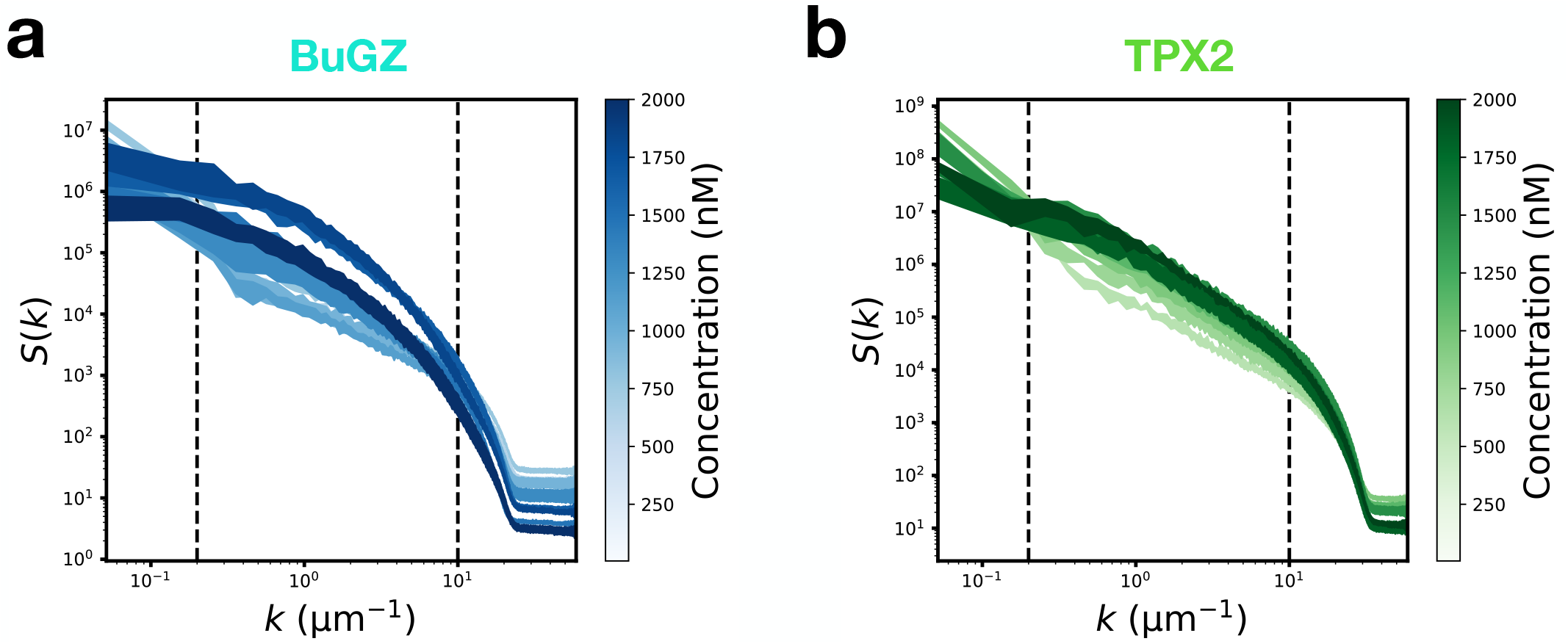
Structure factor of microtubule bundles. Average structure factor *S*(*k*) calculated from the microtubule intensity channels for **a**. BuGZ and **b**. TPX2 as a function of bulk MAP concentration. The set of modes Ω used to compute the order parameter in Fig. 1c-d is demarcated by the black vertical lines. Shaded error bars are standard deviations from *N* = 5 images per concentration.

**FIG. S3.**
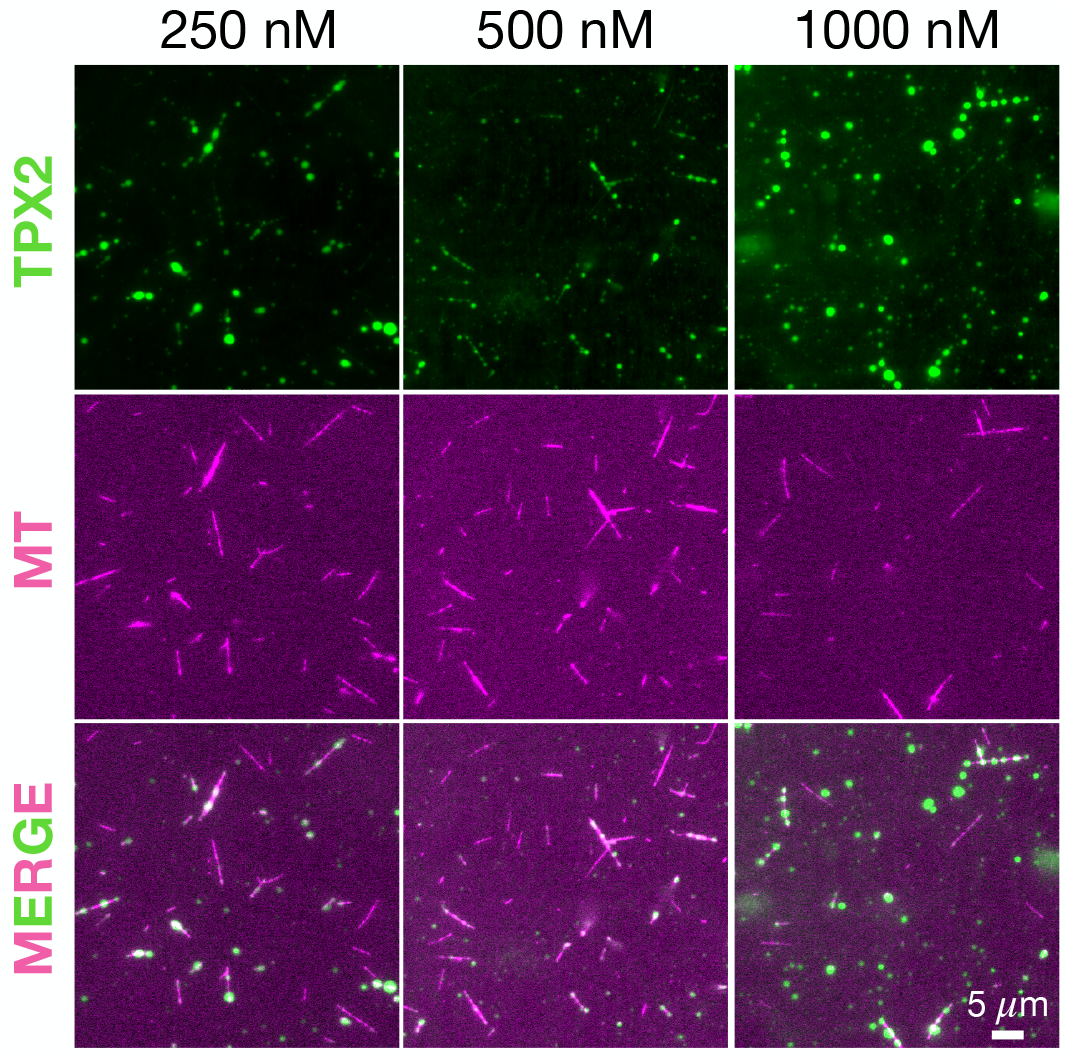
Structure of capillary bundles depend on microtubule density. TIRFM images of microtubule bundles that sedimented to the flow channel surface after a 10 min incubation with TPX2. Experiments done at 1/10th the microtubule density relative to those in Fig. 1. At lower microtubule densities for a given TPX2 concentration, bundles are smaller in size while TPX2 droplets are bigger owing to there being more condensed TPX2 per surface area of microtubules. Scale bar is 5 *µ*m.

**FIG. S4.**
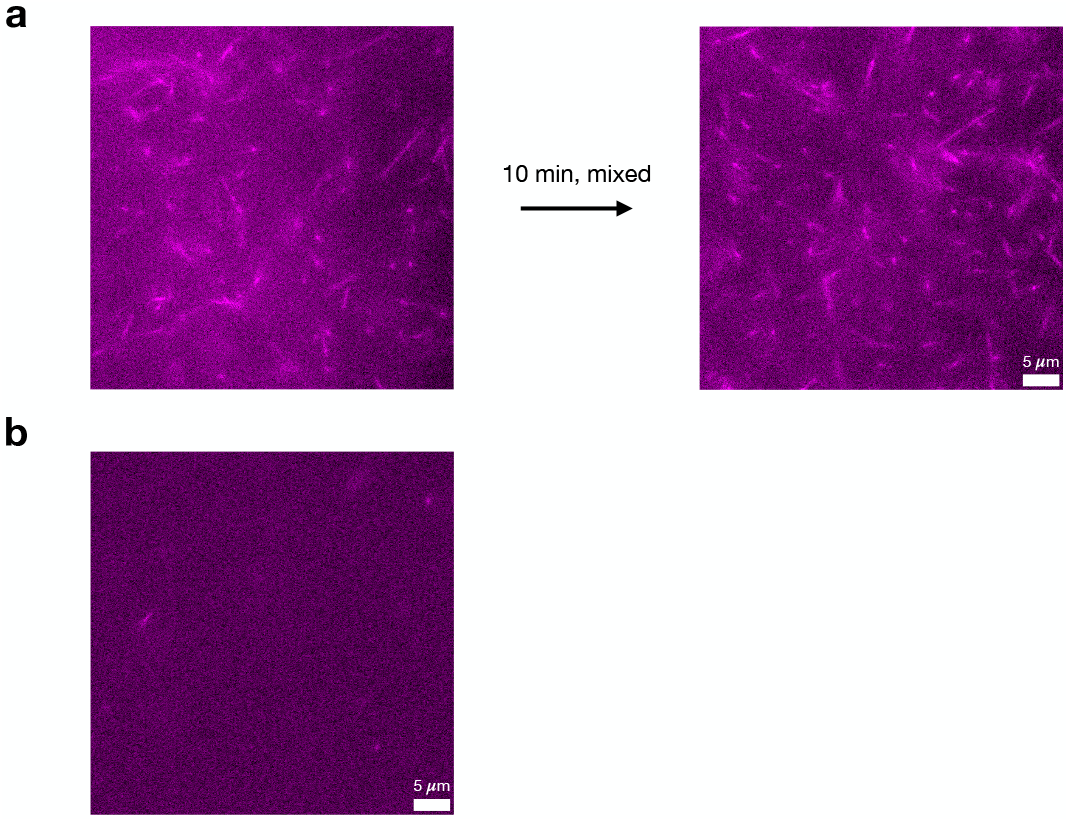
Negative control rules out depletion interaction for microtubule bundling. **a**. Oblique TIRFM images of bulk microtubules before and after mixing in assay buffer without bundling MAPs. This shows that depletion forces alone do not bundle microtubules in our assay buffer and at our tested microtubule densities. **b**. TIRFM images at the surface 10 minutes after mixing show that no microtubule bundles sediment. Scale bars are 5 *µ*m.

**FIG. S5.**
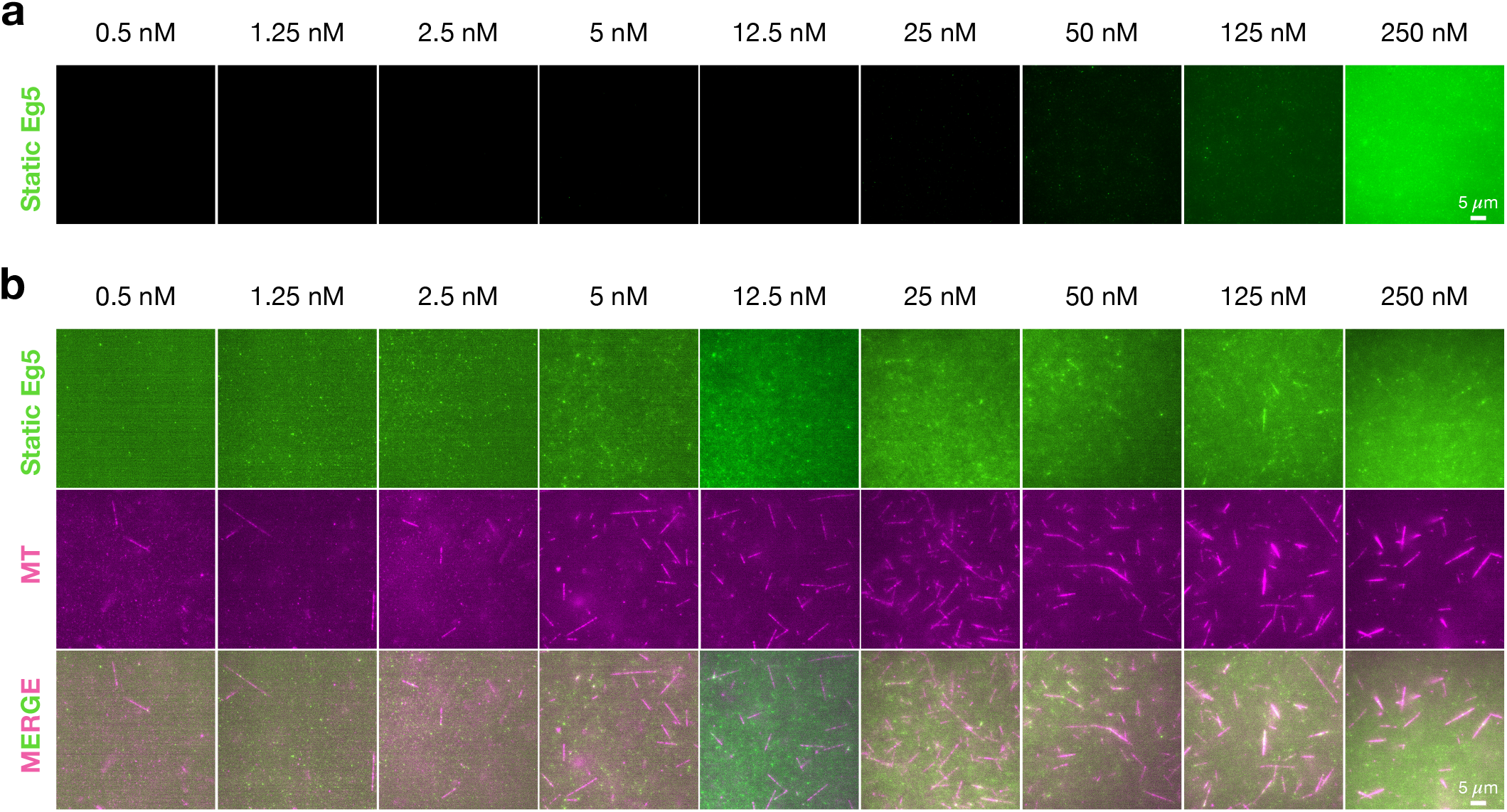
Static Eg5 bulk phase and bundling assay. **a**. Epifluorescence images of bulk phase separation assays for Eg5 at low (1 *µ*M) ATP concentrations (“static Eg5”). No robust mesoscale droplet formation is observed. Lookup tables are the same across concentrations to enable direct comparison. Scale bars are 5 *µ*m. **b**. TIRFM images of microtubule bundles that sedimented to the flow channel surface after a 10 min incubation with static Eg5. Microtubule channel lookup tables are the same across concentrations to enable direct comparison. MAP channel look-up tables are optimized per concentration to allow for visualization. Scale bars are 5 *µ*m.

**FIG. S6.**
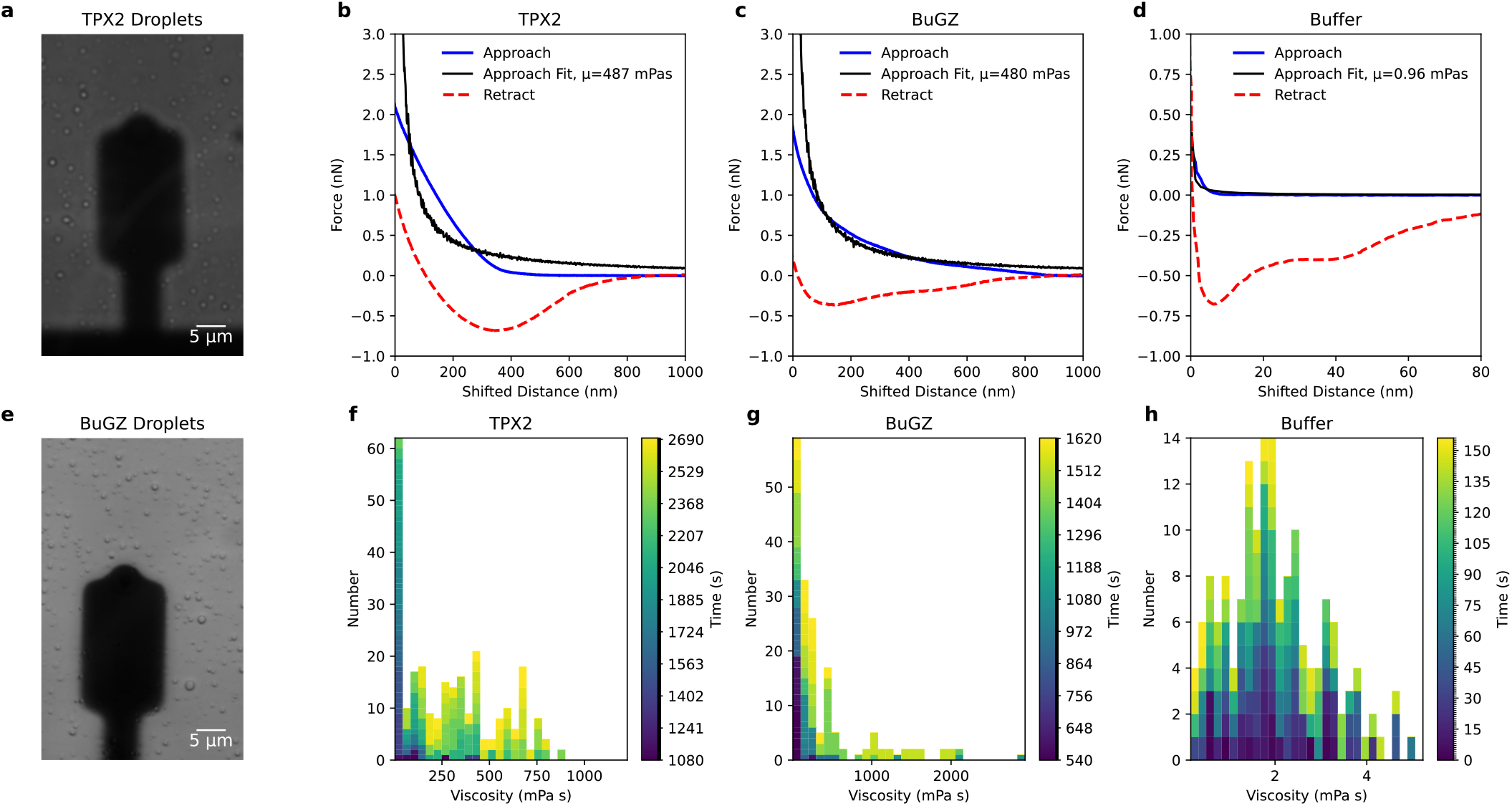
AFM measurements on bulk protein droplets. Brightfield images of droplets below an AFM cantilever for **a**. TPX2 and **e**. BuGZ at 4 *µ*M concentration. Average force curves for **b**. TPX2, **c**. BuGZ, and **d**. buffer including the approach (blue) and retract (red) curves. The shifted distance accounts for the cantilever deflection and estimates the position of the solid glass surface when it reaches zero. The black best fit curve fits the average force curve to the lubrication force prediction of a Newtonian fluid, taking into account the temporally measured average height profile and velocity. The approach fit is used to extract the viscosity, while the retraction minimum for TPX2 and BuGZ are used to estimate the surface tension of the bulk droplet (Supplementary Methods). The fitted viscosities are 487 mPa · s, 480 mPa·s, and 0.96 mPa·s, and the effective inferred surface tensions of the bulk droplets are 50 and 30 *µ*N/m. Histograms of individual fitted viscosities for **f**. TPX2, **g**. BuGZ, and **h**. buffer, with outliers greater than three standard deviations excluded. The histograms include 267, 174, and 163 fitted viscosities from individual approach curves for TPX2, BuGZ, and buffer. The color bar indicates the time at which a given measurement was made after droplets first formed with an error of *±*60 s. The average viscosities from these histograms are 2.0, 303, 383 mPa·s, with standard deviations of 1.1, 257, and 546 mPa·s. 15

**FIG. S7.**
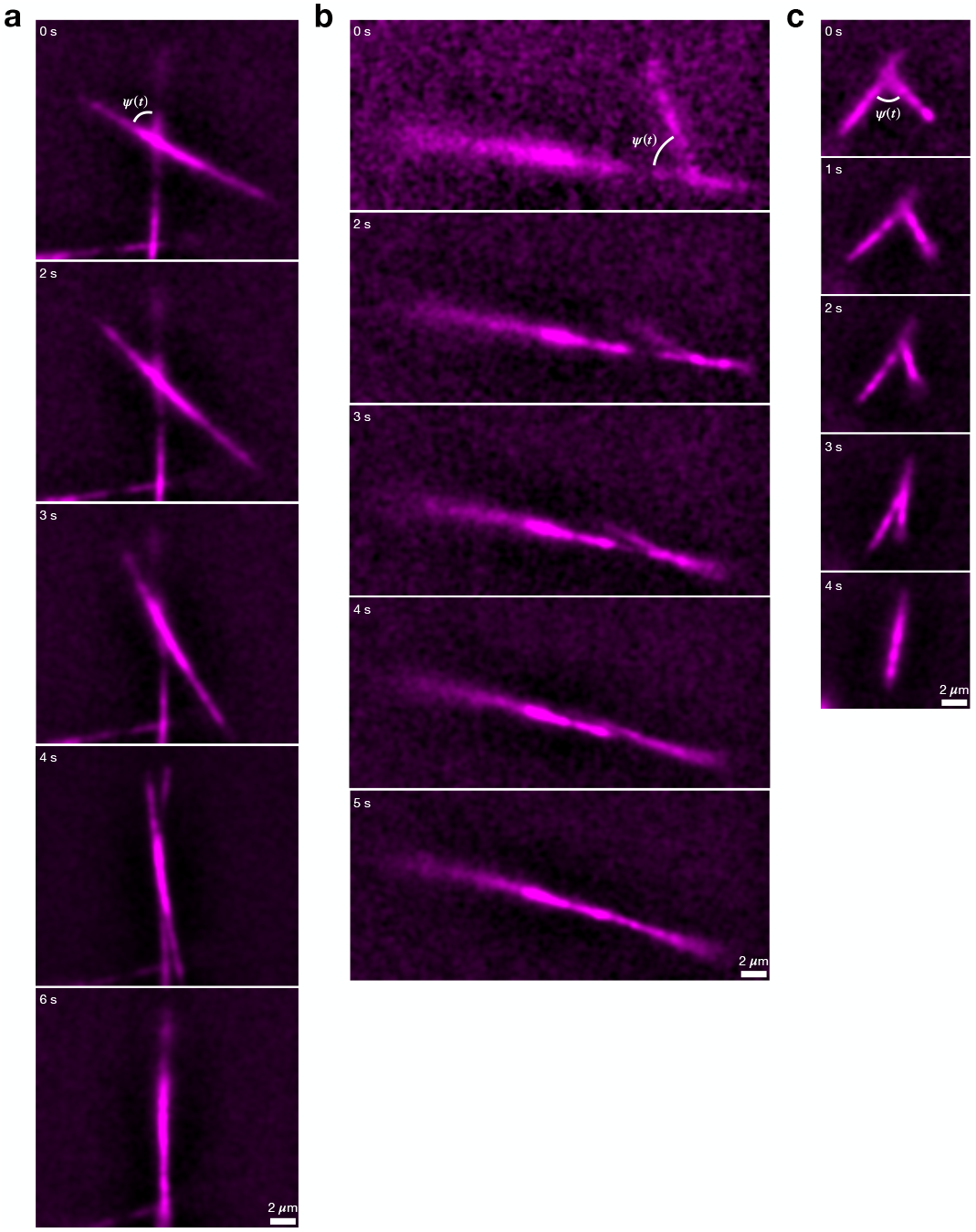
Additional examples of snapping dynamics. Three additional examples of oblique TIRFM live imaging of two bulk microtubule bundles coated with condensed TPX2 above the phase boundary (500 nM) snapping together. Only the microtubule channel is shown. Images were bandpass filtered in Fourier space using a 23 *µ*m lower cutoff and a 310 *µ*m upper cutoff to maximize clarity for visualizing the snapping angle between microtubules. Scale bars are 2 *µ*m.

**FIG. S8.**
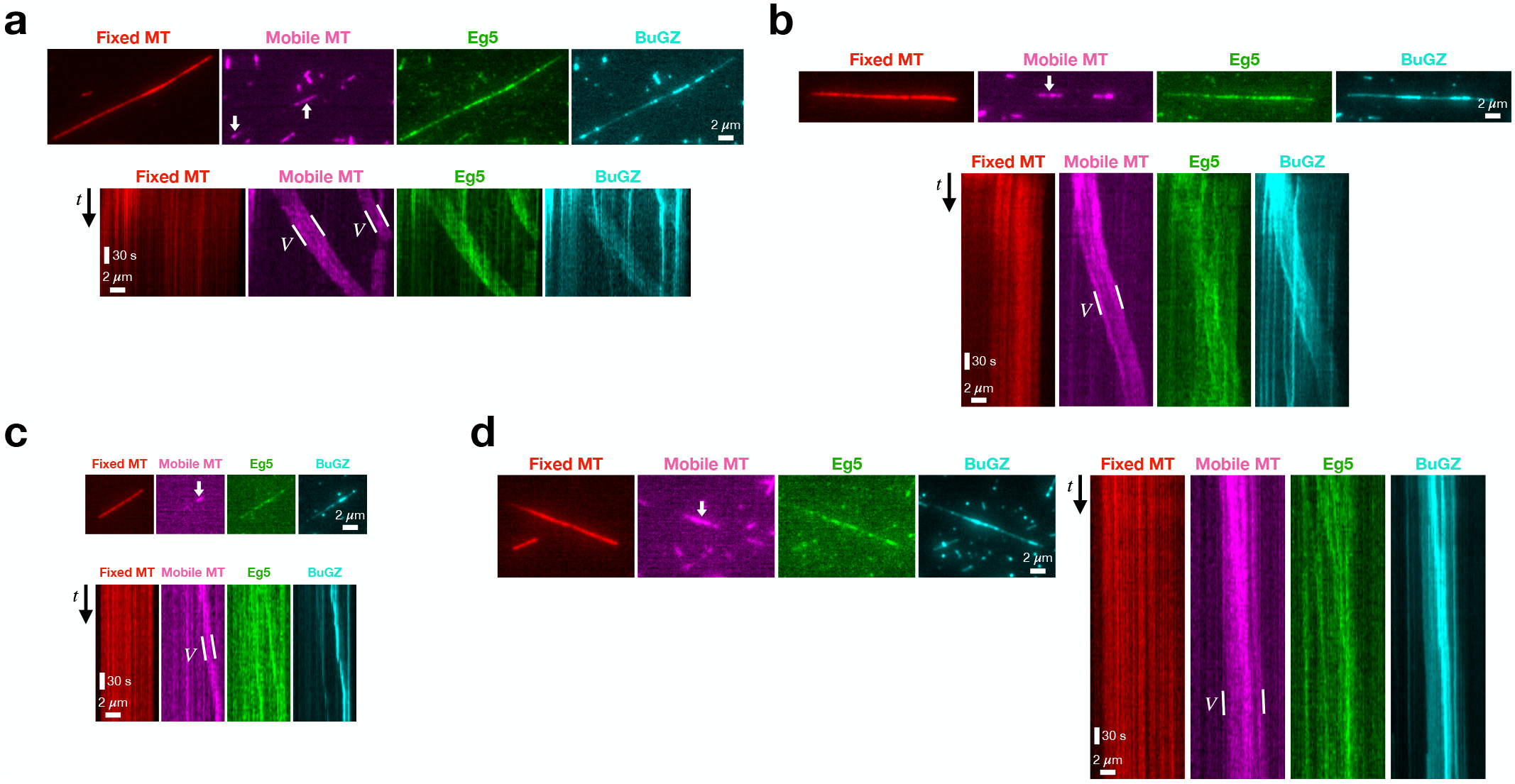
Additional sliding kymographs. Snapshots and kymographs of **a**. 10 nM BuGZ, **b**. 25 nM BuGZ, **c**. 100 nM BuGZ, and **d**. 250 nM BuGZ sliding experiments all at 100 nM Eg5. The velocity *V* of the short, mobile microtubule is computed from the slope of the mobile microtubule kymograph. Scale bars are 2 *µ*m and 30 sec.

**FIG. S9.**
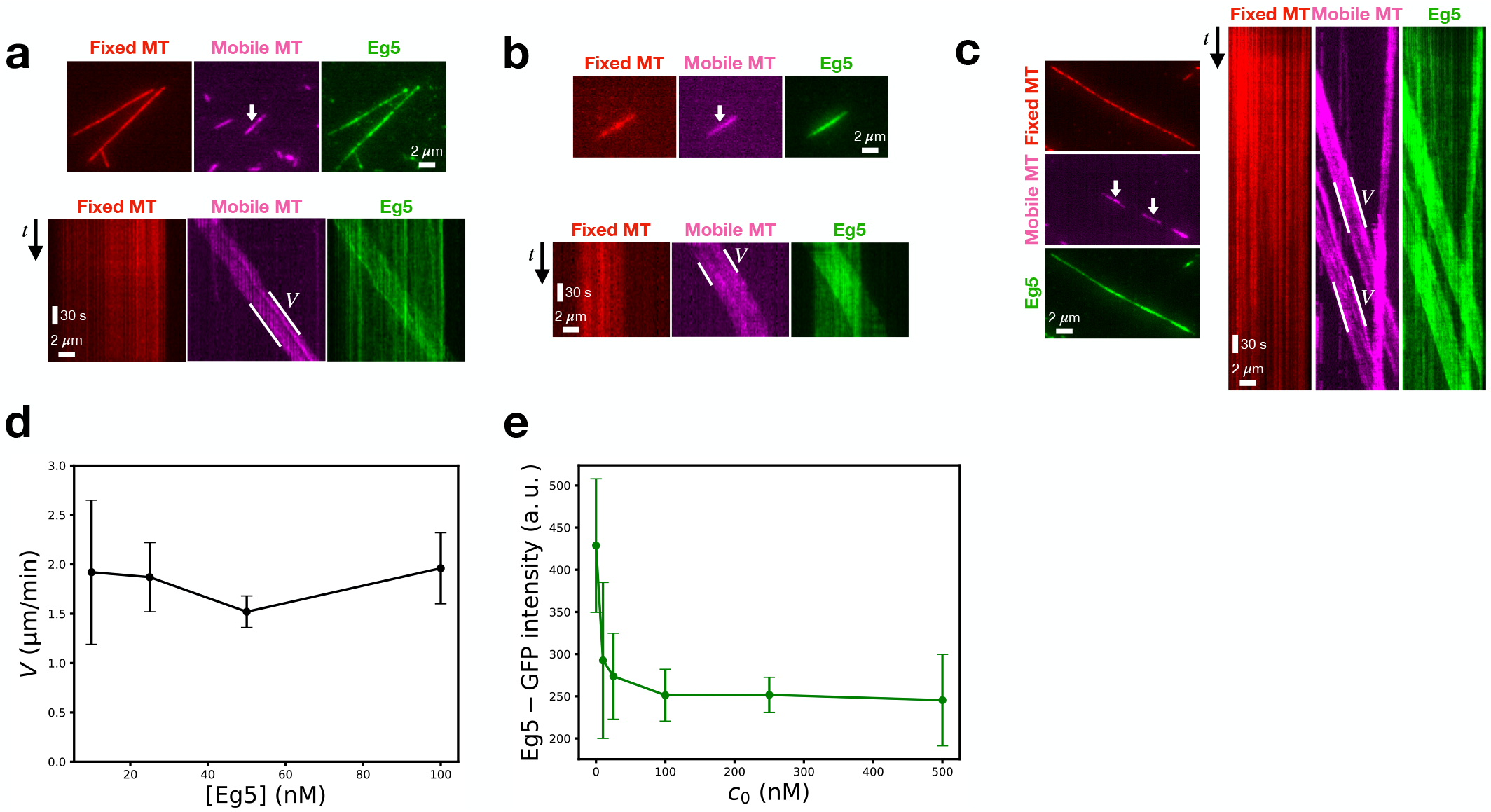
Lower concentration of Eg5 cannot explain observed decrease in sliding velocity. Snapshots and kymographs of **a**. 10 nM Eg5, **b**. 25 nM Eg5, and **c**. 50 nM Eg5 in the absence of BuGZ. The velocity *V* of the short, mobile microtubule is computed from the slope of the mobile microtubule kymograph. Scale bars are 2 *µ*m and 30 sec. **d**. Behavior of the sliding velocity *V* of the short, mobile microtubule as a function of only Eg5 concentration (black line), showing *V* does not measurably depend on the bulk Eg5 concentration. Error bars are standard deviations. *N >* 20 microtubules per condition. **e**. Eg5 fluorescence intensity on microtubules as a function of bulk BuGZ concentration. After measurably falling upon BuGZ addition, Eg5 concentration on microtubules remains roughly constant for all non-zero BuGZ concentrations. Error bars are standard deviations. *N >* 20 microtubules per condition.

## MOVIE LEGENDS

**Movie S1:** Oblique TIRFM live imaging of two bulk microtubule bundles coated with condensed TPX2 above the phase boundary (500 nM) snapping together. Background subtraction was done to enhance visual contrast. TPX2 channel is in green; microtubule channel is in magenta. Scale bar is 5 *µ*m. This movie corresponds to Fig. 3b.

**Movie S2:** Another example of oblique TIRFM live imaging of two bulk microtubule bundles coated with condensed TPX2 above the phase boundary (500 nM) snapping together. Only the microtubule channel is shown. Background subtraction was done to enhance viScale bar is 5 *µ*m. This movie corresponds to Fig. S5a.

**Movie S3:** TIRFM live imaging of Eg5 (green) driven sliding of a mobile short microtubule (magenta) along an anchored long microtubule (red). Background subtraction was done to enhance visual contrast. Scale bar is 1 *µ*m. This movie corresponds to Fig. 4b.

**Movie S4:** TIRFM live imaging of Eg5 (green channel) driven sliding of a mobile short microtubule (magenta) along an anchored long microtubule (red) in the presence of 100 nM BuGZ (cyan). Background subtraction was done to enhance visual contrast. Scale bar is 1 *µ*m. This movie corresponds to Fig. S6c.

## supplementary information

### S1 Theoretical methods

#### S1.1 Capillary force between two microtubules

We first wish to estimate the dynamics at play when two microtubules adhere together under the action of a wetted condensed film with interfacial tension *γ*. Initially, we assume microtubules are adhered in parallel fashion, so that there is only an adhesive force that acts to drive microtubules closer together (Fig. T1a). Quite generally at low Reynolds number, the hydrodynamic drag *F*_*d*_ must balance the adhesive force *F*_*a*_,

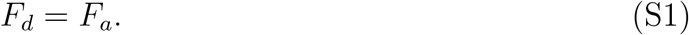

**Figure T1.**
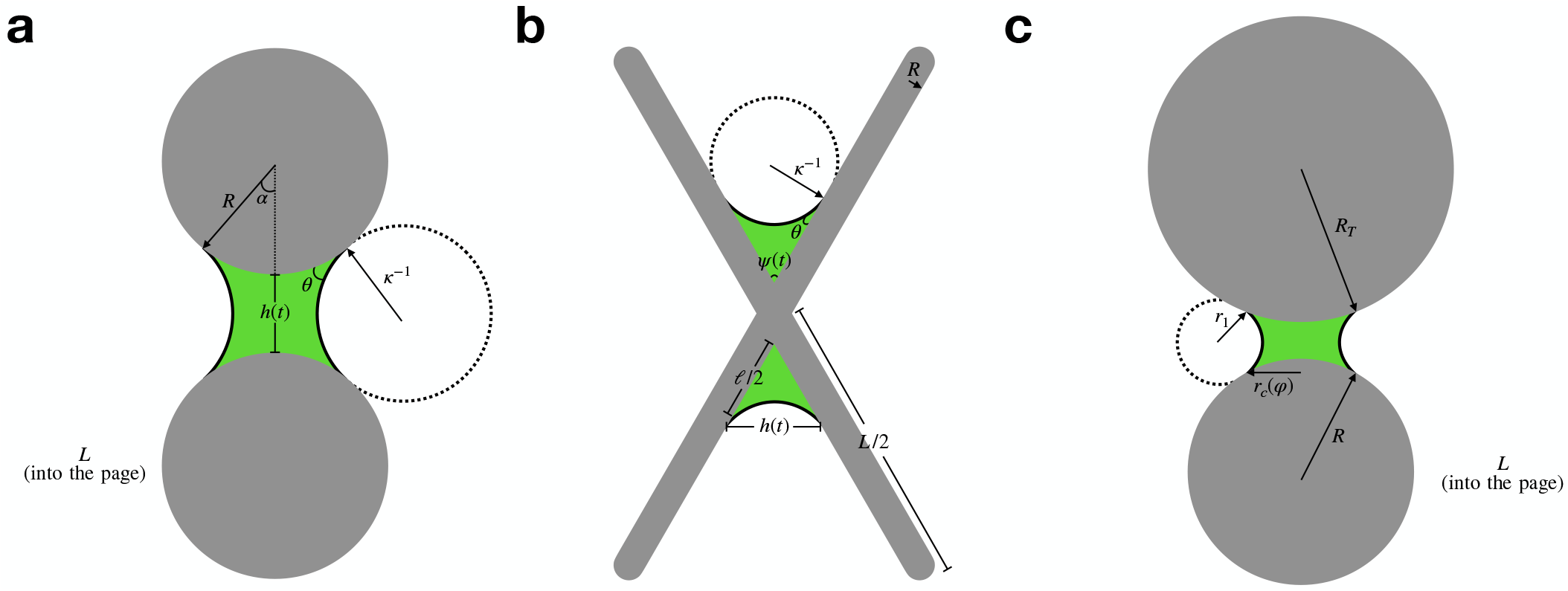
(a) Model for capillary force between two microtubules. Two parallel microtubules of length *L* and radii *R* are adhered together by a capillary bridge of constant cross-sectional area along the length of the microtubule. (b) Model for capillary torque between two microtubules. Two microtubules meet at an acute angle *ψ*(*t*) hinged together by a capillary bridge of wetted length *ℓ*. (c) Model for capillary force between a spherical AFM tip of radius *R*_*T*_ and cylindrical microtubule of radius *R* assuming perfect wetting (*θ* = 0).

To estimate the adhesive force, we assume a condensed film of constant cross section between microtubules of radius *R* and length *L* spaced a distance *h* apart (Fig. T1a). This is a large simplification from the experimental reality, where the condensate morphology is complicated, but it should allow us to capture the basic physics. In so doing, we can use the exact result from Princen [1] to write

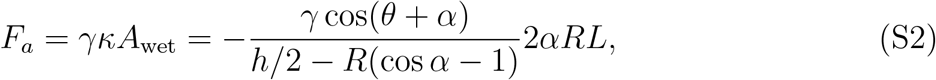

where *κ* is the mean curvature, *A*_wet_ is the wetted area, *θ* is the contact angle, and *α* is a parameter that depends on the volume of the condensed phase in the film, and therefore is set by the bulk concentration of protein.

We consider two possible sources of drag. First is the drag microtubules must overcome to move through the solvent of viscosity *µ*_s_. This is given by [2]

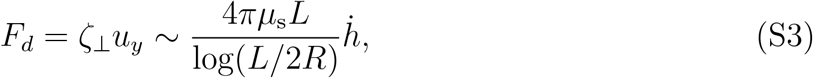

with perpendicular drag coefficient *ζ*_⊥_.

The second source of drag is the flow associated with squeezing the condensed film of viscosity *µ* as the microtubules come together. This squeezing motion mainly generates flows along the microtubule in the horizontal (*x*) direction perpendicular to the direction of squeezing (*y*). Performing a scaling analysis *x* ∼ *b*(*t*), *y* ∼ *h*(*t*), *u*_*x*_ ∼ *U*, and 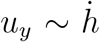 and demanding that continuity ∂_*x*_*u*_*x*_ + ∂_*y*_*u*_*y*_ = 0 hold at O(1) forces

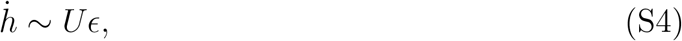

where *ϵ* = *h/b <* 1. We note that the assumption of *ϵ <* 1 gets better over time as the microtubules squeeze together.

Performing the same scaling analysis on the *x* component of the Stokes equations ∂_*x*_*p* = *µ*∇^2^*u*_*x*_ and neglecting terms of O(*ϵ*^2^) forces the pressure *p* to scale like

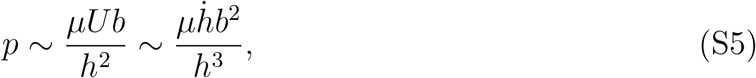

where in the last step we used equation (S4). We note that this squeezing pressure is the dominant resistance to flow as *h* → 0, since the viscous shear stress scales like 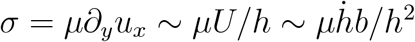.

We now need to relate *b* and *h* to each other using geometry. The profile of the microtubule as measured from the center of the liquid bridge (Fig. T1a) is 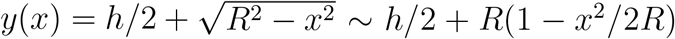, where the last equality is valid if *x < R*, which is an assumption that improves as the squeeze flow progresses. Performing the scaling analysis *x* ∼ *b* we find that if *y* ∼ *h* then *b* must scale like

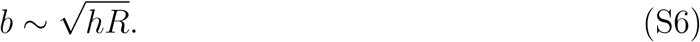

Putting this all together results in the drag force

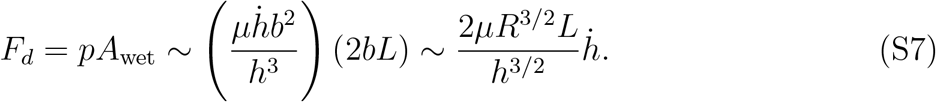

As the squeeze flow progresses and *h* → 0, it is clear that equation (S7) dominates (S3). Thus, equating equation (S7) with equation (S2) results in a separable ODE for *h*(*t*)

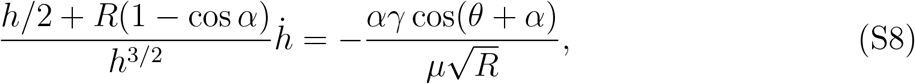

which can be integrated using the initial condition *h*(*t* = 0) = *h*_0_ to give an implicit equation for *h*(*t*)

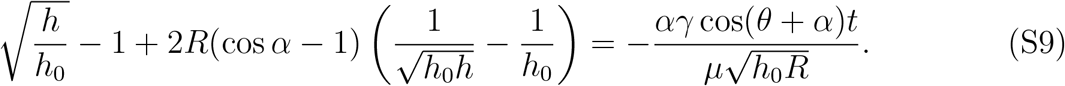

As *h*(*t* → ∞) → 0, there is a dominant balance between only two terms in equation (S9) which results in the final asymptotic scaling

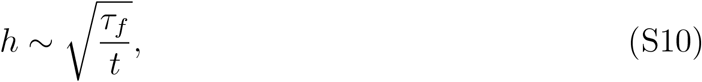

where the timescale 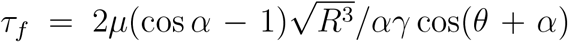 emerges as the dominant capillary time that governs the squeezing dynamics. This is to say that, under action of capillary forces alone, the microtubules will get arbitrarily close to each other with a spacing between them that decreases as 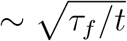. Of course, this theory breaks down at the length scale of single molecules, at which point a more complicated disjoining pressure arises that likely results in a finite equilibrium separation [3].

#### S1.2 Capillary torque between two microtubules

We now move to consider the rotational dynamics at play when two microtubules contact each other at an acute angle *ψ* (Fig. T1b). At low Reynolds number, the adhesive capillary torque *T*_*a*_ must balance the resistive viscous torque *T*_*d*_,

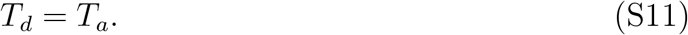

An exact calculation of *T*_*a*_ is complex and depends on the intricate details of the highly contorted interface between angled filaments [4, 5]. However, to make analytical progress we argue that the capillary pressure that drives filaments to snap together is dominated by the in-plane radius of curvature (Fig. T1b). The out-of-plane radius of curvature serves principally to decrease the out-of-plane distance *h* between filaments, which was already considered in Sec. (S1.1) (Fig. T1a). Because *L* ≫ *h*, it is reasonable to carry out the calculation entirely in the 2d plane. Using plane geometry (Fig. T1b), the radius of curvature is therefore *κ*^−1^ = −*ℓ* sin(*ψ/*2)*/*2 cos(*ψ/*2 + *θ*), where *ℓ* is the wetted length of the droplet on the filaments that drives the snapping dynamics. This gives rise to the capillary force

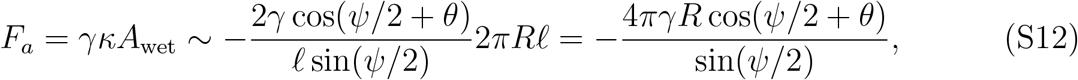

where for simplicity we assume the condensate wets the entire circumference of the microtubule. The corresponding torque is

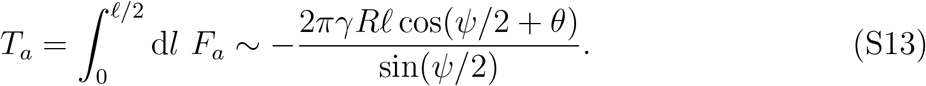

For the viscous torque, we must consider two dynamical regimes. The first regime is where *ψ*(*t*) is close to the initial angle *ψ*_0_, which can be large and finite as observed in our experiments. In this case, the wetted length *ℓ* satisfies *ℓ* ≪ *L* and can be treated as a constant *ℓ* = *ℓ*_0_ throughout the motion. The resistance to snapping is dominated by rotating microtubules of length *L* through the solvent of viscosity *µ*_s_ (Fig. T1b), which is given by [2]

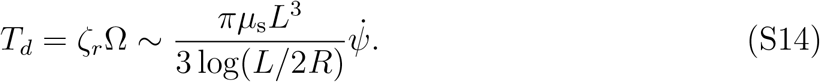

Equating equations (S13) and (S14) results in the ODE for *ψ*(*t*)

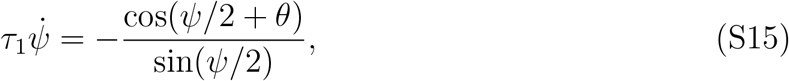

where the timescale that governs the early time snapping dynamics falls out as 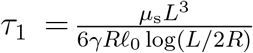. We can integrate this ODE to give an implicit equation for *ψ*(*t*). Taking *ψ*(*t* = 0) = *ψ*_0_, we find

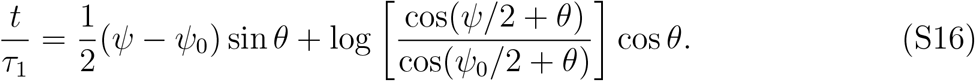

If we assuming perfect wetting *θ* = 0, a reasonable assumption for condensed MAPs on microtubules, we find the following explicit solution for *ψ*(*t*),

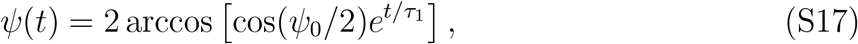

which can be expanded for early times *t* → 0 dropping terms of O(*t*^2^) and higher to find

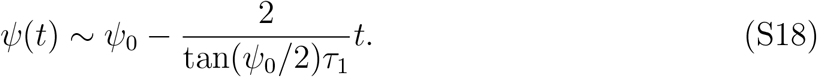

Even though equation (S18) was derived in the *ψ*(*t* → 0) → *ψ*_0_ limit, this linear function can represent most of the experimentally measured data in Fig. 2c well, and predicts that microtubules will take a time *T*_1_ = (*ψ*_0_*/*2) tan(*ψ*_0_*/*2)*τ*_1_ to fully snap together.

The second regime is where *ψ*(*t*) → 0, when the microtubules are snapping together and have to squeeze the condensate of viscosity *µ* between the aligning microtubules (Fig. T1b). In this case, *ℓ* = *ℓ*(*t*) as the wetted length moves significantly along the microtubules during the squeeze flow. Taking *h*(*t*) as the horizontal distance between snapping filaments (Fig. T1b), the same arguments that resulted in equation (S5) can be applied here, such that the dominant viscous resistance is the squeezing pressure which scales like

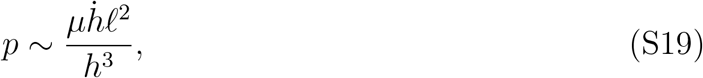

so long as *h < ℓ*, an assumption that improves as the squeeze flow progresses as *ψ* → 0. To make progress, we relate *h* to *ψ* and *ℓ* using the law of cosines

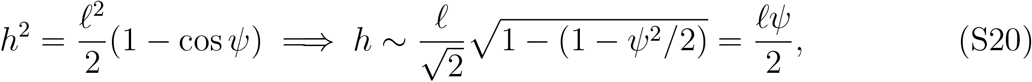

where the second relation is valid for small *ψ*. Therefore 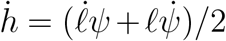. We can further relate *ℓ* and *ψ* by employing the constraint that the condensate volume remains constant during the motion. In this 2D calculation, we can approximately represent this by the triangular area *A* enclosed by *h* and the two wetted lengths *ℓ/*2. Computing the area of the condensate exactly would result in an unwieldy elliptic integral that is not needed to model the dominant physics of *h* decreasing as *ℓ* increases during the snapping dynamics. The simple triangular geometry therefore results in *A* = (*ℓ*^2^*/*8) sin *ψ* ∼ (*ℓ*^2^*/*8)*ψ* for small *ψ*. Therefore

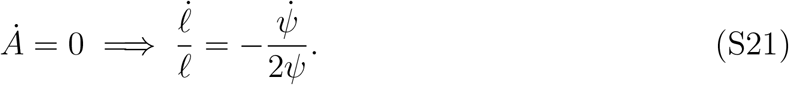

Substituting this into the expression for 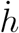 gives 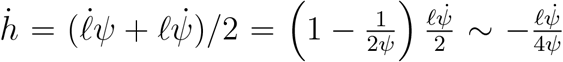 for small *ψ*. Substituting these results into equation gives

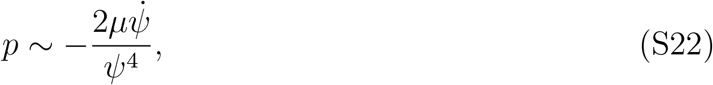

and therefore the resulting force is

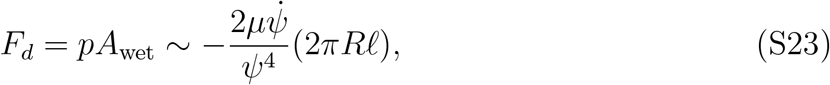

from which the viscous torque may be computed as

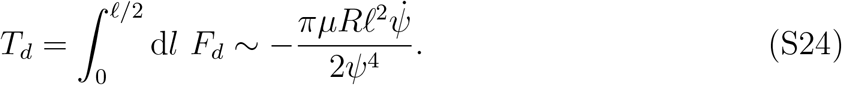

We can expand the capillary torque in equation (S13) for *ψ* → 0 to give

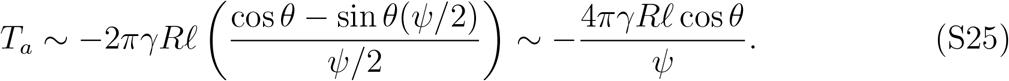

Equating equations (S24) and (S25) gives us our final ODE for *ψ*(*t*)

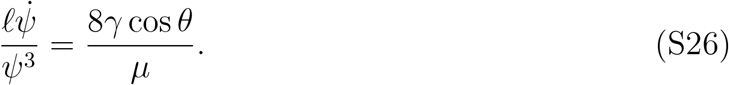

At this stage it is most convenient to eliminate *ℓ* in favor of *ψ* by integrating the geometric constraint equation (S21). Taking *ψ*(*t* = 0) = *ψ*_0_ and *ℓ*(*t* = 0) = *ℓ*_0_, we have

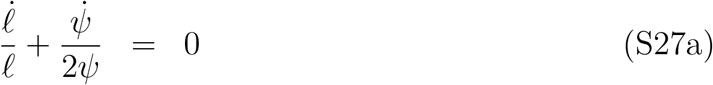

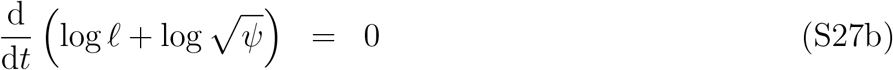

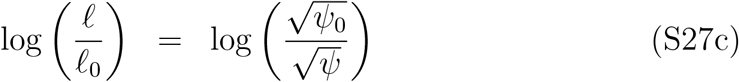

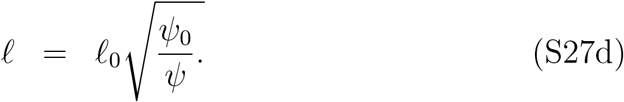

Substituting this result into equation (S26) gives

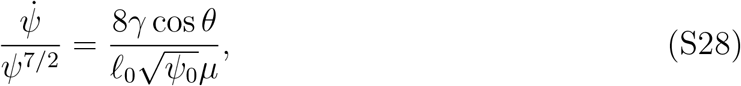

which can be directly integrated to give

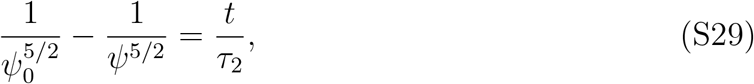

where 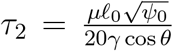 is the timescale that governs the late time snapping dynamics. As *ψ*(*t* → ∞) → 0 a dominant balances emerges resulting in the final asymptotic scaling

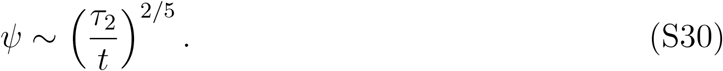

While eventually we expect this lubrication scaling to dominate for late times, we do not have the experimental spatiotemporal resolution to observe the relavant small angle dynamics in this work.

#### S1.3 Comparing mesoscale capillary bridges to single-molecule crosslinkers

To more solidly contrast snapping mechanisms of crosslinking and capillary forces, the dynamics based on crosslinking can be analyzed more precisely. If the crosslinkers act as effective springs that connect when the distance between opposing microtubules is less than 2*b* and supply a spring force per cross linker of *f*_*cl*_ = − *kx*, where *x* is the distance between neighboring microtubule surfaces, *x* = *l* sin(*ψ/*2), with *l* describing a coordinate parallel with the microtubule, and *k* a spring constant. With a crosslinking density per unit length, *q*, the force in the *x* direction would be

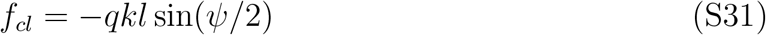

The adhesive torque would then be:

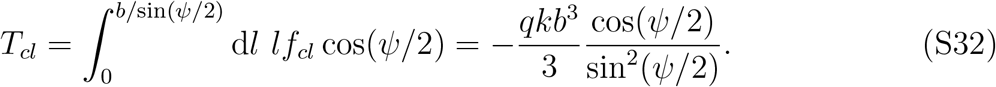

Here, *b/*sin(*ψ/*2) describes the distance over which the two crossed microtubules will be connected by crosslinkers.

Again, we can balance the adhesive torque with the torque required to rotate a microtubule of length *L, T*_*d*_. Since here there is no viscous condensate driving the adhesion, we assume that the dominant viscous torque is the one needed to rotate the microtubule in the solvent of viscosity *µ*_*s*_,

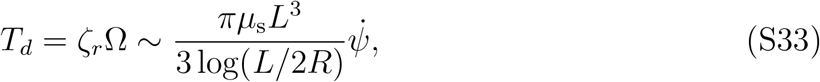

to arrive at the differential equation governing snapping:

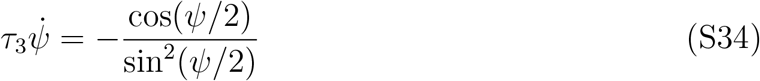

where 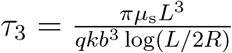

We are interested in the dynamics of *ψ*(*t*) between *ψ*(*t* = 0) = *ψ*_0_ and when the entirety of the microtbule is connected, *ψ* = 2 sin^−1^(2*R*_0_*/L*). Here, we will assume that *R*_0_ ≪ *L*.

The solution to the differential equation gives an implicit expression for *ψ*(*t*)

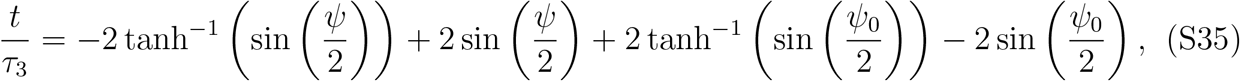

which if *ψ* is always is small, can be simplified to:

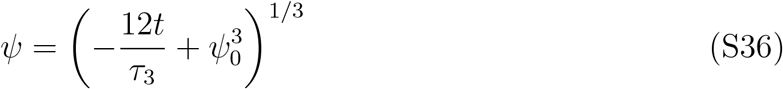

This time course function shape appears quite similar to the time course predicted for the first regime of capillary snapping. From the shape of the snapping curves, it is difficult to differentiate directly between mechanisms of capillary snapping driven on a time scale *τ*_1_ and *τ*_3_.

We can instead compare the different expectations for the timescales of snapping based on ratios of these effective time scales, for example:

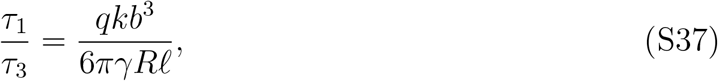

where we have now substituted *ℓ*_0_ → *ℓ*. If we assume similar adhesion force scales between the two models so *kb* ≈ *γR*, and we take *qb* ≈ 1, we find that the ratio of time scales is sensitively governed by the ratio *b/ℓ. b* is always a molecular scale, whereas *ℓ* is at the scale of the wetting film, which can reach the order of the microtubule length *L* for late times, *b* ≪ *ℓ*. Because b is a molecular length scale and l is the mesoscopic length scale of the wetted film, we expect *τ*_1_ ≪ *τ*_3_. That is, even for comparable adhesion forces, we generally expect capillary torques to be at least an order of magnitude larger than crosslinking torques due to their mesoscale nature. This means that the crosslinking dynamics are expected to act much more slowly than the capillary snapping, although the system-specific details will determine by how much.

#### S1.4 Capillary force between a spherical tip and a microtubule

Here, we estimate the force between a sphere (to represent the AFM tip) and a cylinder (to represent the microtubule) with a connecting capillary bridge within the Derjaguin approximation [6] and assuming perfect wetting. We assume that the sphere of radius *R*_*T*_ is in contact with a cylinder with radius *R*, which we define as the origin of our coordinate system. A condensate bridge with constant radius of curvature *r*_1_ perfectly wets the two surfaces. The Laplace pressure will be approximately:

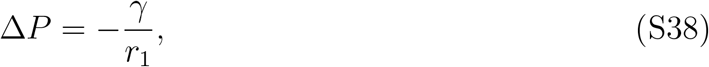

as long as the contact area is small relative to the radii of the confining surfaces and the contact line at *r*_*c*_(*φ*) is much larger than *r*_1_. At equilibrium, the radius of curvature and contact angles will be fixed regardless of the azimuthal angle from the origin.

We can define the bottom surface of a sphere with a function,

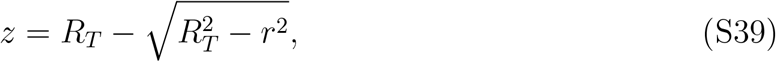

and similarly for the cylinder

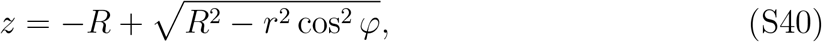

where *r* and *φ* are given by cylindrical coordinates centered at the origin. If we assume that the capillary bridge is small relative to *R*_*T*_ and *R* at leading order, *r* ≪ *R*_*T*_, *R*, then the distance between the two surfaces can be approximated as

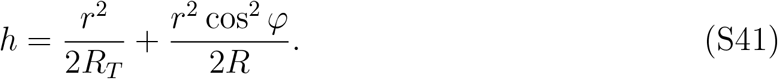

Based on the perfect wetting assumption, we have *h* = 2*r*_1_. Suppose that the contact line exists at some angular-dependent position *r*_*c*_(*φ*). In order for the system to be at equilibrium, the value of *r*_1_ must be fixed at every angle. From this requirement, we get a definition for *r*_*c*_(*φ*)

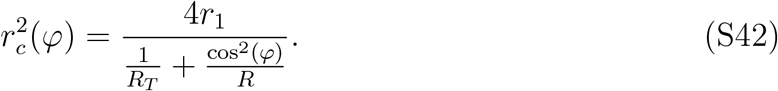

In order to find the capillary force, we need to compute the product of the Laplace pressure multiplied by the area of contact. The area of contact, *A*, can be calculated by direct integration,

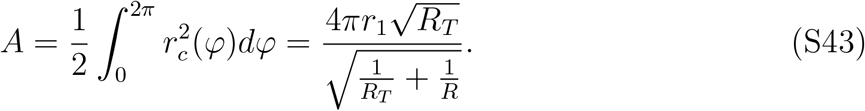

Therefore, the capillary force at contact, *F*, can be approximated as:

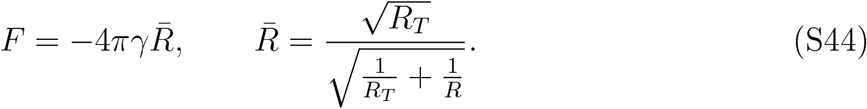

In the limit of *R* → ∞ the expression for a sphere interacting with a plane is recovered. In the limit of *R*_*T*_ → ∞, the force diverges since the capillary bridge extends over the entire (infinite) cylinder.

#### S1.5 Phase field model of the condensate interface

##### S1.5.1 Flory-Huggins Cahn-Hilliard Model

Here, we employ the standard mean-field Flory-Huggins [7, 8] model to represent the protein (biopolymer) thermodynamics, and we combine the homogeneous part with a Cahn-Hilliard [9] term to capture the surface energy. The dimensionlesss free energy density 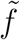, at temperature *T*, monomer and solvent volume *v*, and with Boltzmann constant *k*_*B*_ is

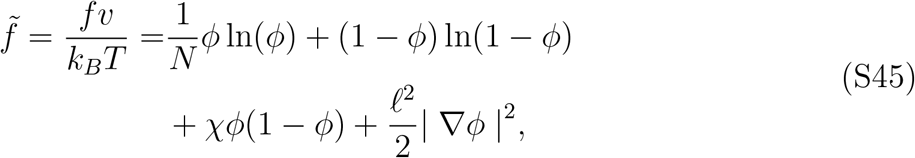

where *ϕ* is the volume fraction of the protein, *N* is its length, *χ* is the Flory interaction parameter, and *ℓ* is a length scale that characterizes the phase boundary thickness that is on the scale of an individual monomer. From this point forward, the gradients of composition will be assumed to only occur in one coordinate direction, *x*, such that | ∇*ϕ*|^2^= |*ϕ*^*′*^|^2^.

In order to derive an analytical expression for the surface energy, we will expand the free energy density around its critical point and truncate at a few terms. The critical point, where *f* ^*′′′*^(*ϕ*) = 0, is defined at a composition of *ϕ*_*c*_ and interaction strength *χ*_*c*_,

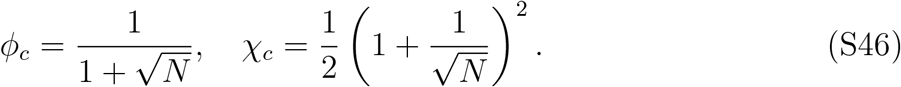

The free energy density expanded around the critical point up to fourth order in *δϕ* = *ϕ* − *ϕ*_*c*_ and neglecting terms that are constant or linear in *ϕ* gives

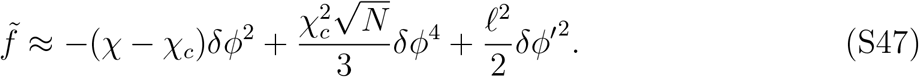

We can derive a generalized chemical potential that includes contributions from the interface by taking a variational derivative, 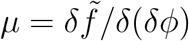,

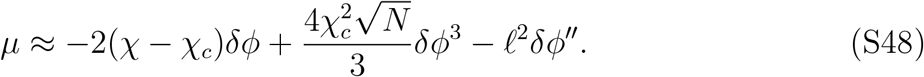

Near the critical point, the homogeneous part of the chemical potential is an odd function. Chemical equilibrium between the dilute and concentrated phase implies that: *δϕ*_+_ = −*δϕ*_−_. Therefore, we have in the bulk phases near the critical point [10],

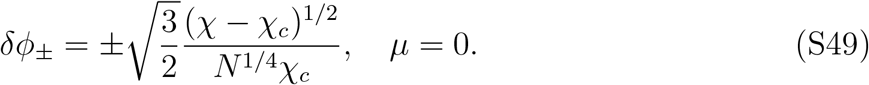

Next, we solve for the 1D profile across the phase boundary between composition *δϕ*_−_ as *x* → −∞ and composition *δϕ*_+_ as *x* → + ∞, relative to the interface at *x* = 0, by applying *µ* = 0 at all values of *x*. We define the following rescalings into dimensionless variables 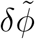 and 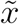 to simplify the calculations,

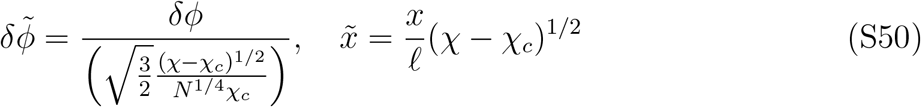

to render the equilibrium chemical potential equation as:

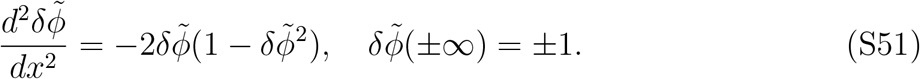

The solution to the above equation is:

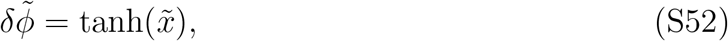

or with dimensions now returned

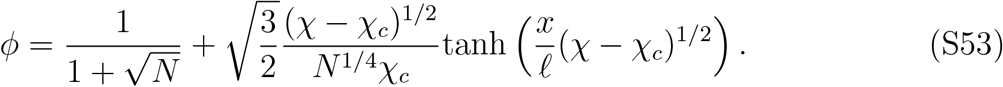

The surface tension can be derived by integrating the excess free energy over the interface region [9],

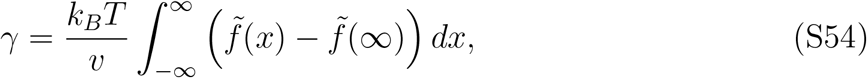

or in in other terms

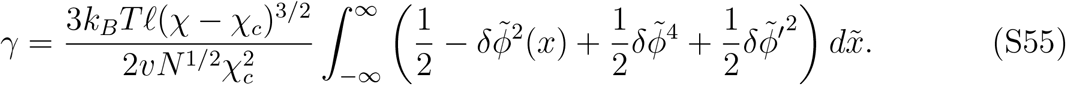

We can evaluate the integral analytically to get a final expression for the surface tension in the mean-field limit,

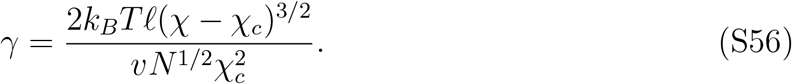

The above expression highlights that the surface tension increases with increasing interaction strength, *χ*, but decreases as the protein sequence length, *N*, increases. Next, we rationalize the possible trends in *χ* in the context of electrostatic interactions, similar to ref. [11], since charge interactions can strongly influence phase separation and thus surface tension of condensates.

##### S1.5.2 Estimating the electrostatic contribution to the free energy density

In what follows, we will derive the leading-order contribution of electrostatic interactions to the Flory interaction parameter, *χ*, and thereby connect the protein charge interactions to the surface tension of their condensed phases. The Voorn-Overbeek [12, 13] model combines the Flory-Huggins model of polymer entropy with a Debye-Huckel type free energy density from the screening by ions in an electrolyte. The electrostatic contribution to the free energy density is given by

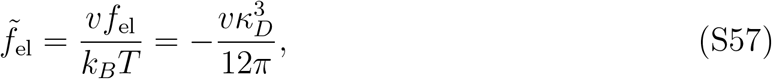

where *κ*_*D*_ is the inverse Debye length. The protein is treated as an unconnected polyelectrolyte, such that it also contributes to the ionic strength of the solution. Therefore, the inverse Debye length is defined as,

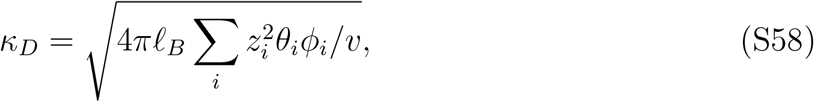

where the summation over species *i* corresponds to all species, including the salt ions and protein. *ℓ*_*B*_ is the Bjerrum length which describes the length at which two point charges interact with energy *k*_*B*_*T*, defined as

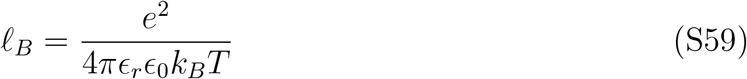

where *e* is an elementary charge, *ϵ*_*r*_ is the dielectric permittivity of the solvent, and *ϵ*_0_ is the permittivity of free space.

Here, we are assuming that all monomers and salt ions occupy the same volume, *v. θ*_*i*_ corresponds to the total fraction of charged sites on each species (the sum of both the number of positive and negative sites), where we assume the salt ions are completely dissociated *θ*_*±*_ = 1. Further, we assume that all charged sites on the protein carry a charge of *z*_*i*_ *±* 1.

While the model is able to capture basic trends in the salt-dependent phase separation of polyelectrolytes and their electrostatic interactions, it is not a perfect model for concentrated electrolytes. Further, it neglects the correlations in the charge sequences along the protein [14] and structuring in the electrolyte [13]. Therefore, we use this model to provide general relationships with respect to protein charge, but we do not expect it to be quantitatively accurate for all model systems.

The free energy density can be recast in the convenient form,

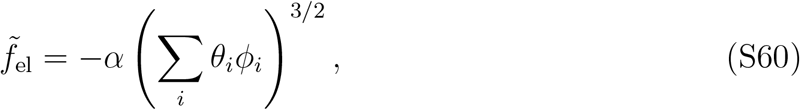

where the parameter *α* is defined by

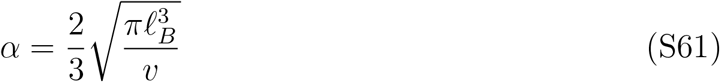

To find the leading order electrostatic contributions to the free energy density, we can expand the result in the limit of small protein volume fraction. If we expand the free energy density around a protein concentration of *ϕ*_*p*_ = 0, we get the Flory-Huggins model with a salt dependent *χ* contribution, *χ*_el_.

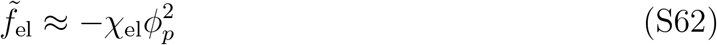

where:

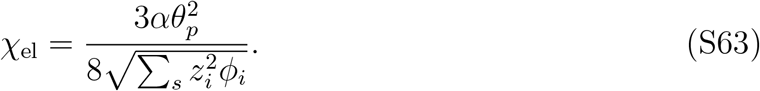

The summation in the denominator is now only over the salt ion species. Here, we see that the leading order electrostatic contribution to the Flory interaction parameter goes with the square of protein charge and decreases inversely proportional to the square root of salt concentration. Note that the actual value of *χ* may include some non-electrostatic contribution, *χ*_0_, so that

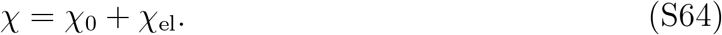

To compare BuGZ and TPX2, the total fraction of charged residues, *θ*_*p*_, were estimated from the amino acid sequences using the EMBOSS pK values [15, 16] for the charged residues at pH 6.8 of the BRB80 buffer. For TPX2, *θ*_*p*_ = 0.33 and for BuGZ, *θ*_*p*_ = 0.20. ^1^

In the physiological solutions at room temperature, *ℓ*_*B*_ = 0.7 nm, and we can roughly assume that *ℓ* = 0.3 nm and *v* = (0.3nm)^3^ to characterize the size of solvent, monomers, and salt ions. With these parameters, the value of *α* is 4.2. The BRB80 electrolyte has an ionic strength of about 160 mM. Therefore, the values of *χ*_el_ for TPX2 and BuGZ, respectively, are 2.3 and 0.9. Note that these predicted values are quite sensitive to the chosen parameters *ℓ* and *v*, although their ratio will not change significantly.

To estimate the expected difference in surface tension between TPX2 and BuGZ, we only need to define *N* based on the sequence length. The values of *N* for TPX2 and BuGZ are 694 and 959 respectively. The mean-field theory therefore would predict that from electrostatic interactions alone, the surface tension of TPX2 and BuGZ should roughly differ by a factor of 10, which is what we observed experimentally. Even so, the meanfield theory overpredicts the absolute value of *γ* by two orders of magnitude, highlighting the possible inaccuracies introduced through the expansions and truncations around the critical point.

#### S1.6 Sliding resistance from a viscous film

Here we consider the hydrodynamic resistance experienced by parallel microtubules sliding apart due to a constant force *F*. In our experiments, this force is provided by Eg5 motors sliding one mobile microtubule relative to another that is chemically fixed to the surface.

In the case where there is just solvent of viscosity *µ*_*s*_ between the microtubules, the resistance to sliding is dominated by the classic slender-body result 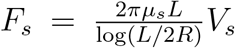, where *V*_*s*_ is the measured sliding speed in this case.

In the case where there is an additional condensed film of viscosity *µ* between the microtubules, the motion is resisted by both the slender-body term as well as the viscous film of thickness *h*. The simplest approximation for this additional resistance that captures the basic features is 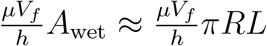, and therefore 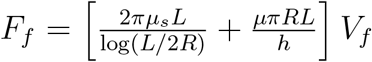, where *V*_*f*_ is the measured sliding speed in this case.

Assuming the Eg5 driving force is the same in both scenarios, we can set *F*_*s*_ = *F*_*f*_ and compute the ratio *V*_*f*_ */V*_*s*_, which serves as a prediction for how much the condensed film slows down the Eg5-driven sliding,

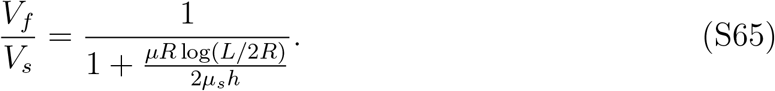

While these are rather simple scaling arguments, we believe they capture the basic physics. The real drag expression in the presence of the condensed film results from a complicated multiphase flow problem, and since *R < h* simplifying lubrication approximations do not apply. Such an effort would be outside the scope of this work

### S2 Experimental methods

#### S2.1 Protein expression and purification

Full-length TPX2 (*X. laevis*, Gene ID: 398174) with a N-terminal StrepII-6xHis-GFPTEV tag was cloned into a pST50 vector, transformed into Rosetta2 *E. Coli* cells, grown up to an OD_600_ ≈ 0.6 at 37^*□*^C in 2 L of LB broth, and then expressed using 0.75 mM IPTG at 25^*□*^C for 7 hr. Cells were pelleted, homogenized, then lysed using an Emulsiflex in lysis buffer (50 mM Tris-HCl, 750 mM NaCl, 15 mM imidazole, 6 mM BME, pH 8.0) containing 2.5 mM PMSF, 10 *µ*g/ml DNAse I, and 2 cOmplete EDTA-free protease inhibitor tablet. Lysate was centrifuged at 30,000 RPM on a 45 Ti rotor for 30 min and the supernatant was bound to Ni-NTA agarose beads equilibrated in lysis buffer. Protein was eluted with lysis buffer containing 200 mM imidazole and then further purified using a Superdex 200 HiLoad 16/600 SEC column into storage buffer (10 mM HEPES, 500 mM KCl, 1 mM MgCl2, 5 mM EGTA, 10% w/v sucrose, pH 7.7).

Full-length BuGZ (*H. sapiens*, Gene ID: 7756) with a C-terminal StrepII-6xHis-BFP-TEV tag was cloned into a pST50 vector, transformed into Rosetta2 *E. Coli* cells, grown up to an OD_600_ 0.6 at 37^*□*^C in 2 L of LB broth, and then expressed using ≈ 0.60 mM IPTG at 16^*□*^C for 16 hr. Cells were pelleted, homogenized, then lysed using an Emulsiflex in lysis buffer (50 mM Tris-HCl, 500 mM KCl, 1 mM MgCl2, 1 mM DTT, pH 7.8) containing 200 *µ*M PMSF, 10 *µ*g/ml DNAse I, and 2 cOmplete EDTA-free protease inhibitor tablet. Lysate was centrifuged at 30,000 RPM on a 45 Ti rotor for 30 min and the supernatant was bound to CV = 5 mL of equilibrated Strep-Tactin resin in lysis buffer. After a 1 hr incubation on a shaker, the mixture was transferred to a column and washed with 10 CV of lysis buffer. Protein was eluted with 3 CV of lysis buffer containing 2.5 mM desthiobiotin and then further purified using a Superdex 75 10/300 SEC column into storage buffer (25 mM HEPES, 500 mM KCl, 1 mM MgCl2, 1 mM DTT, 15% v/v glycerol, pH 7.7).

Full-length kinesin-5 (Eg5) ((*X. laevis*, Gene ID: 379112)) with a C-terminal StrepII-6xHis-GFP-TEV tag was cloned into a pFastBac vector which was used to infect 1 L of *S. frugiperda* (Sf9) cells for a 2 day expression. Virus-infected cells were pelleted then lysed using an Emulsiflex in lysis buffer (50 mM HEPES, 300 mM KCl, 1 mM MgCl2, 1 mM DTT, 0.2 mM Mg-ATP, pH 8.0) containing 200 *µ*M PMSF, 10 *mu*g/ml DNAse I, and 1 cOmplete EDTA-free protease inhibitor tablet. Lysate was centrifuged at 50,000 RPM on a 70 Ti rotor for 50 min and the supernatant was bound to CV = 2 mL of equilibrated Strep-Tactin resin in lysis buffer. After a 1 hr incubation on a shaker, the mixture was transferred to a column and washed with 10 CV of lysis buffer. Protein was eluted with 3 CV of lysis buffer containing 2.5 mM desthiobiotin and then further purified using a Superose 6 10/300 SEC column into storage buffer (25 mM HEPES, 500 mM KCl, 1 mM MgCl2, 1 mM DTT, 15% v/v glycerol, pH 7.7).

All purified recombinant proteins were flash frozen in working aliquots using liquid nitrogen and stored at −80^*□*^C.

#### S2.2 Stabilized microtubule seed preparation

To make short, double-cycled, GMPCPP-stabilized, Atto-647 labelled microtubule seeds, a 40 *µ*L mixture of 20 *µ*M bovine tubulin (10% labelled with Atto-647 dye) and 1 mM GMPCPP in BRB80 (80 mM PIPES, 1 mM MgCl2, 1 mM EGTA, pH 6.8) was polymerized for 45 min in a 37^*□*^C water bath and centrifuged at 126000 g for 8 min at 30^*□*^C in a TLA-100 rotor. The supernatant was discarded and the seeds were resuspended in cold BRB80 and allowed to depolymerize on ice for 20 min. GMPCPP was then added up to 1 mM and the mixture was polymerized again for 45 min in a 37^*□*^C water bath and centrifuged at 126000 g for 8 min at 30^*□*^C in a TLA-100 rotor. The supernatant was discarded and the seeds were resuspended in room temperature BRB80. The seeds were pipetted into 1 *µ*L aliquots, flash frozen in liquid nitrogen, and stored at 80− ^*□*^C.

To make long, GMPCPP-stabilized, biotinylated, Alexa-568 labelled microtubule seeds, a 40 *µ*L mixture of 20 *µ*M bovine tubulin (10% labelled with Alexa-568 dye, 10% labelled with biotin) and 1 mM GMPCPP in BRB80 was polymerized for 2 hr in a 37^*□*^C water bath and centrifuged at 13000 RPM for 8 min at room temperature in a tabletop centrifuge. The supernatant was discarded and the seeds were resuspended with room temperature BRB80 containing 1 mM GMPCPP. The seeds were incubated overnight in the dark at room temperature and were used within 2-3 days.

#### S2.3 *In vitro* bulk condensation assay

Purified protein (either GFP-TPX2 or BuGZ-BFP) was precleared of aggregates by centrifugation at 80000 RPM for 10 min at 4^*□*^C in a TLA-100 rotor. The supernatant was diluted to a target concentration in assay buffer (25 mM HEPES, 25 mM KCl (low salt) or 100 mM KCl (high salt), 1 mM MgCl2, 50 *µ*g/ml *κ*-casein, pH 7.7), mixed thoroughly, and pipetted into a flow chamber. The flow chamber was sealed with nail polish and incubated coverslip side down for 10 minutes to allow condensates to settle. Condensates were imaged using epifluorescence on a Nikon Ti-E microscope with a 100x objective and 1.49 numerical aperture. Exposure times and LED power were consistent across all tested 16 concentrations for each protein. An ORCA-Fusion BT digital CMOS camera was used for acquisition.

Phase diagrams were computed as the area fraction of the condensed phase versus bulk concentration. The condensed phase was defined as regions having an intensity value above the threshold calculated using Otsu’s method [17] for the highest concentration images for each protein.

#### S2.4 *In vitro* bundling assay

Double-cycled, Atto-647 microtubule seeds were diluted in assay buffer (25 mM HEPES, 100 mM KCl, 1 mM MgCl2, 50 *µ*g/ml *κ*-casein, pH 7.7) and mixed thoroughly. Purified protein (either GFP-TPX2 or BuGZ-BFP) was precleared of aggregates by centrifugation at 80000 RPM for 10 min at 4^*□*^C in a TLA-100 rotor. The supernatant was added up to a final target protein concentration and a final seed dilution of 1*/*100 or 1*/*1000. The mixture was mixed twice with a p20 pipette and pipetted into a flow chamber. The flow chamber was sealed with nail polish and incubated coverslip side down for 10 minutes to allow bundles to settle. Bundles were imaged using 2-color TIRF. Exposure times and laser power were consistent across all tested concentrations for each protein. While the bundles are imaged on the glass coverslip, they form in the bulk region of the flow chamber before they settle to the glass surface.

For live snapping assays, coverslip-bottomed culture well plates were washed three times with assay buffer. lMicrotubule seeds were added, mixed thoroughly, and then imaged using oblique TIRF, well above the plane of the coverslip to avoid surface interactions. Once a good field of view was established, acquisition was started on the 647 channel only to maximize frame rates. Precleared GFP-TPX2 was pipetted into the well up to a final 500 nM concentration and a final seed dilution of 1*/*100 and mixed once with a p20 pipette. Acquisition was stopped after 5-10 min. Microtubule lengths were estimated using the straight line tool in Fiji (ImageJ) on the frame when the microtubules were best in focus.

All assays were done on a Nikon Ti-E microscope with a 100x objective and 1.49 numerical aperture. An ORCA-Fusion BT digital CMOS camera was used for acquisition. The average structure factor *S*(*k*) was calculated from the microtubule intensity channels for each MAP concentration. For each image *I*(*x, y*), the power spectrum *P* (*k*_*x*_, *k*_*y*_) = |ℱ [*I*(*x, y*)] |^2^ is computed, where ℱ is the two-dimensional Fourier transform. *P* (*k*_*x*_, *k*_*y*_) is converted to polar coordinates *P* (*k, θ*) and then *S*(*k*) is calculated via 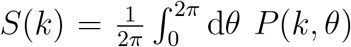. The order parameter is then defined as 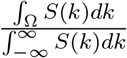, which represents the relative power of the modes Ω that characterize the bundling transition (Fig. S2).

#### S2.5 *In vitro* atomic force microscopy assay on microtubules

Short, GMPCPP-stabilized microtubule seeds were diluted 1*/*10 in BRB80 and electrostatically adhered to an atomically smooth mica surface pre-incubated with a 1 M MgCl_2_ solution for 10 min [18]. Unbound seeds were washed off with fresh BRB80, and the final solution with the desired protein concentration was pipetted onto the mica surface. Samples were probed with a silicon-nitride AFM cantilever tip (MLCT-BIO DC Bruker) with a nominal radius of 20 nm. Since the silicon nitride tip is negatively charged at the pH of BRB80 [19], analogous to the microtubule lattice, we assume that condensates which wet microtubules would similarly wet the AFM tip. Using the JPK NanoWizard 4 atomic force microscope, we first localized microtubules by generating elevation maps using tapping mode (Peakforce QNM). Then, using force spectroscopy mode, we performed a series of slow force ramps at specific locations along the length of a chosen microtubule in order to measure interfacial forces directly on the microtubule lattice. The force measurement ramps were carried out at velocity of 0.5 *µ*m/s over 0.1 *µ*m, with a force threshold of 0.2 nN.

The force curves were processed as follows. All force curves were shifted by a constant value equal to the average force between 98 and 100 nm from the surface so that the baseline was at a force of 0 pN. Some force curves (not plotted) were rejected if their baseline was not flat, judged by applying a cutoff of 0.02 nN difference between 90 nm and 100 nm. If the difference in the baseline far from the surface exceeded this value, it was assumed that the beginning of the ramp was not sufficiently quiescent for reliable force measurements. No filters or other smoothing of the raw data was applied. The value of the adhesion force for each force ramp was calculated by taking the minimum of the force curve. To calculate the effective surface tension, the minima of the average curves was used to extract an average adhesion force at contact. The adhesion energy was calculated by integration of the force over distance. In order to minimize errors from integrating over the baseline regions where noise might bias the integral, a linear subtraction fitted to the data between 60 and 100 nm was performed from the cumulative integral. The adhesion energy was calculated as the minimum of this corrected cumulative integral.

#### S2.6 *In vitro* atomic force microscopy assay on bulk droplets

To supplement the microtubule-specific measurements that quantify capillary forces directly on the microtubule lattice, measurements are made directly on large (∼ *µ*m scale) condensate droplets sedimented onto a glass substrate to estimate condensate/water surface tension, *γ*, and the condensate viscosity, *µ*.

The experiment begins at *t* = 0 when condensate are diluted from a high salt storage buffer with BRB80 buffer to arrive at a final concentration of 4 *µ*M in a glass bottomed petri dish (uncoated, ibidi *µ*-Dish 35 mm, low). Using a Zeiss LSM 900 / Axio Observer 7 with a long focal distance air objective (Objective LD PN 63x/0.75 Korr Ph2), the DirectOverlay functionality is used to co-localize the AFM tip (Bruker SAA-SPH-1UM, with spring constant *k* ≈ 0.25 N/m and radius *R* = 1*µ*m) and the droplets. Ten force ramp measurements are made on an individual droplet, and the process is repeated for 18 and 15 droplets for TPX2 and BuGZ, respectively. The force ramp is initiated on the first droplet once a droplet of sufficient size has sedimented within the microscope field of view and the cantilever position can be calibrated with DirectOverlay, about 18 minutes for TPX2 and 9 minutes for BuGZ. Between each droplet measurement, the DirectOverlay calibration is repeated to ensure proper alignment with the sedimented droplet of choice. Droplets are chosen at random within the field of view. Example images of droplets at the end of the experiment are shown for TPX2 and BuGZ in Fig. S6 (a,e). We note that due to the small droplet sizes, the contact angle of the droplets on the glass substrates were not able to be detected reliably.

The force ramp is conducted at a fixed piezo velocity of 10 *µ*m/s with a ramp height of 4 *µ*m up to a force threshhold of 4 nN. From the approach curve, one can fit the effective viscosity from the repulsive force upon approach to the rigid glass interface, properly accounting for the cantilever deflection. Here, the corrected distance between the cantilever tip and the solid surface, *H*, is computed by correcting for the cantilever deflection as estimated by the contact slope of the measured force, *F* (*x*). We define *x* as the piezo displacement coordinate, with *x* = 0 defined as the position where the force reaches its maximum, *F*_0_. Following [20], if *m* is the absolute value of the contact slope (considered to be in the compliant zone and representative of the effective spring constant of the cantilever tip in contact with the surface), the contact position *x*_*c*_ is estimated as *x*_*c*_ = *F*_0_*/m* and the cantilever deflection at each position is given as *F* (*x*)*/m*. Therefore, the value of H can be represented as

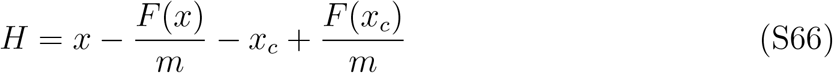

from this *H* function, the velocity *dH/dt* can be computed directly from the time series data.

To extract viscosity, we use the lubrication forces of a sphere near a plane [20],

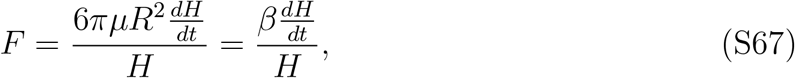

valid when *H* ≪ *R*. We restrict the fit to the region in which the distance of the sphere to the glass plane is less than 500 nm (0 *< H <* 500 nm) or the closest point where the force is greater than 10 pN (*F >* 10 pN), so that the lubrication approximation is a decent approximation and the force magnitude is above noise.

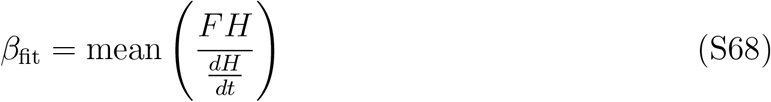

where the fit is taken only over the allowable range of *H* and *F* specified above.

The baselines are subtracted by the force value at *x* = 200 nm for buffer and *x* = 2.0 *µ*m for the condensates TPX2 and BuGZ. Force curves are filtered out by assessing the drift in the baseline at *x* = 2.45 − 2.5 *µ*m if the average force in this region is greater than 0.5 nN after baseline subtraction.

This procedure is repeated for each individual approach force curve to create a histogram in Fig. S6 (f-h), rejecting only fits that are greater than three standard deviations away from the mean. In the histogram, the segments of the bars are colored by the time at which each force ramp was initiated, with *t* = 0 corresponding to when the condensate is first formed via dilution (time errors for *t* = 0 could be at most 60 seconds from the estimate, but all other time shifts are extracted directly from metadata). The fitting procedure is also done on the averaged curves in Fig. S6 (b-d). First, the force curves averaged at each *x*, then the fitting procedure, including calculating *H*, is done for that averaged force curve.

For the control conditions of buffer without condensate, we see that in Fig. S6 (c), the viscosity of a water is effectively recovered. All measurements are made around a temperature of *T* = 24^*□*^C, where the viscosity of water is 0.91 mPas. The measured viscosity is averaged out to be 0.96 mPas (legend). While the average force curve closely reproduces the value of water, the histogram of individually fitted force curves illustrates the spread across force ramps. Due to the noise in the force curves, there is a distribution of values of the viscosity for different measurements, although the trends in the fitted viscosity do not appear to be a function of time, as would be expected for a simple fluid containing only water and salt. The actual viscosity of water is within one standard of deviation (1.1 mPas) of the average value extracted from all force curves (2.0 mPas).

For BuGZ and TPX2, on the other hand, we get a much larger fitted viscosity than water, within the range of viscosities reported for condensates but on the lower end. The viscosity fitted to the average force curves are 487 mPas and 480 mPas for TPX2 and BuGZ, respectively. For TPX2, the shape of the average force curve does not closely model that of the lubrication approximation, likely indicating that there are elastic contributions to the response due to the condensates viscoelasticity. On the other hand, BuGZ is much better behaved, and appears to have a force curve more consistent with the lubrication analysis assuming a Newtonian liquid. The histogram averages are for 303 mPas and 383 mPas, with standard deviation of 257 mPas and 546 mPas, for TPX2 and BuGZ, respectively. There is a clear trend in both condensate systems that at later times, larger viscosity values are measured, corresponding to droplet aging.

Next, we describe the procedure used for estimating the surface tension of the condensed droplets, *γ*, that have been sedimented onto glass. At contact, for a small, perfectly wetting droplet between a sphere in contact with a flat interface, the adhesive force is:

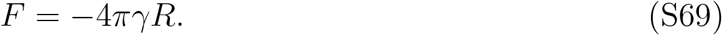

We estimate the adhesive forces from the retraction force curve minimums for TPX2 and 20 BuGZ. Note there is an adhesive minimum also for the case of a buffer without condensate present, but this minimum occurs in close contact with the surface. On the other hand, the apparent distance of the capillary adhesion driven by condensate appears at larger effective distances from the interface (*H >* 100 nm). While the fact that the observed capillary adhesion minimums do not occur precisely at contact may call into question the validity of the above contact expression, the force minimum on retraction can nevertheless be used to give a rough estimate of the surface tension within the context of a contact capillary bridge between glass and the cantilever tip. For TPX2 and BuGZ, respectively, the force minimums are −0.65 nN and −0.37 nN, respectively. These correspond to a surface tension of *γ*_bulk_ ≈ 50 *µ*N/m and *γ*_bulk_ ≈ 30 *µ*N/m, respectively. While the BuGZ value is in perfect agreement, we measure about a factor of 4 lower interfacial tension for bulk TPX2 droplets compared to condensed TPX2 on microtubules. Therefore, in the case of condensed TPX2, subtle nanoscale effects are at play in setting the value of the interfacial tension on microtubules, such as interactions with C-terminal tubulin tails. Other discrepancies in the value for TPX2 may indicate non-idealities in the droplet geometry compared to the simplistic model, to non-equilibrium effects in the measurements, or to the unknown effective contact angles of the condensate wetting interfaces. Because of the droplets’ small size, it is difficult to determine the contact angle they make with the glass surface.

#### S2.7 *In vitro* motor sliding assay

Silanized and biotinylated coverslips were prepared for this assay. Coverslips were sonicated in 3 M NaOH for 30 minutes, washed with MilliQ water, then sonicated in Piranha solution for 45 minutes. Coverslips were then washed with MilliQ water and spin dried. 2-3 drops of GOPTS (3-glycidyloxypropyl trimethoxysilane) were sandwiched between two coverslips and placed in a 75^*□*^C oven for 30 min. Coverslips were then separated, washed with acetone, then dried with nitrogen gas. 1 g of HO-PEG-NH2 and 100 mg of biotin-CONH-PEG-NH2 were mixed, sandwiched between two coverslips, and incuvated overnight in a 75^*□*^C oven. The next day coverslips were separated, sonicated in MilliQ water for 30 min, washed with MilliQ water, then spin dried. Coverslips were stored at 4^*□*^C and used within 1-2 months.

A flow channel was made using a silanzed and biotinylated coverslip and incubated with 5% Pluronic F-127 solution for 5 min. The channel was washed with BRB80 (80 mM PIPES, 1 mM MgCl2, 1 mM EGTA, pH 6.8) and then incubated with 0.1 mg/ml NeutrAvidin for 10 min. The channel was washed with BRB80 then incubated with long Alexa-568 biotin microtubule seeds diluted 1*/*500 in BRB80 for 10 minutes. The channel was washed with assay buffer (25 mM HEPES, 25 mM KCl, 1 mM MgCl2, pH 7.5; 1% v/v glycerol, 50*µ*g/ml *κ*-casein, 10 mM BME, 1 *µ*M Mg-ATP, 20 mM glucose) then incubated with 100 nM of Eg5-GFP for 1 min. The channel was washed with assay buffer then incubated with short Atto-647 microtubule seeds diluted 1*/*200 in assay buffer for 3 min. The channel was washed with assay buffer, the final reaction mixture (assay buffer containing target BuGZ-BFP concentration, 100 nM Eg5-GFP, 5 mM Mg-ATP, 320 *µ*g/ml glucose oxidase, 55 *µ*g/ml catalase) was pipetted in, then the channel was sealed with nail polish. The reaction was imaged using 4-color TIRF on a Nikon Ti-E microscope with a 100x objective and 1.49 numerical aperture. An ORCA-Fusion BT digital CMOS camera or an Andor Zyla scientific CMOS camera was used for acquisition targeting around 2 seconds per frame.

## Footnotes

1 In the main text, the symbol *f* is used in place of *θ*_*p*_ to avoid confusion with the contact angle, *θ*.

